# Tyrosine kinase inhibitors induce mitochondrial dysfunction during cardiomyocyte differentiation through alteration of GATA4-mediated networks

**DOI:** 10.1101/2020.05.04.077024

**Authors:** Qing Liu, Haodi Wu, Qing-Jun Luo, Chao Jiang, Zhana Duren, Kevin Van Bortle, Ming-tao Zhao, Bingqing Zhao, Jun Liu, David P Marciano, Brittany Lee-McMullen, Chenchen Zhu, Anil M Narasimha, Joshua J Gruber, Andrew M Lipchik, Hongchao Guo, Nathaniel K Watson, Ming-Shian Tsai, Takaaki Furihata, Lei Tian, Eric Wei, Yingxin Li, Lars M Steinmetz, Wing Hung Wong, Mark A. Kay, Joseph C Wu, Michael P Snyder

## Abstract

Maternal drug exposure during pregnancy increases the risks of developmental cardiotoxicity, leading to congenital heart defects (CHDs). In this study, we used human stem cells as an *in-vitro* system to interrogate the mechanisms underlying drug-induced toxicity during cardiomyocyte differentiation, including anticancer tyrosine kinase inhibitor (TKI) drugs (imatinib, sunitinib, and vandetanib). H1-ESCs were treated with these drugs at sublethal levels during cardiomyocyte differentiation. We found that early exposure to TKIs during differentiation induced obvious toxic effects in differentiated cardiomyocytes, including disarranged sarcomere structure, interrupted Ca^2+^-handling, and impaired mitochondrial function. As sunitinib exposure showed the most significant developmental cardiotoxicity of all TKIs, we further examine its effect with in-vivo experiments. Maternal sunitinib exposure caused fetal death, bioaccumulation, and histopathologic changes in the neonatal mice. Integrative analysis of both transcriptomic and chromatin accessibility landscapes revealed that TKI-exposure altered GATA4-mediated regulatory network, which included key mitochondrial genes. Overexpression of GATA4 with CRISPR-activation restored morphologies, contraction, and mitochondria function in cardiomyocytes upon TKI exposure early during differentiation. Altogether, our study identified a novel crosstalk mechanism between GATA4 activity and mitochondrial function during cardiomyocyte differentiation, and revealed potential therapeutic approaches for reducing TKI-induced developmental cardiotoxicity for human health.

**Highlights:**

- Early-stage exposure to TKIs induced cardiotoxicity and mitochondrial dysfunction
- GATA4 transcriptional activity is inhibited by TKIs
- Network analysis reveals interactions between GATA4 and mitochondrial genes
- GATA4-overexpression rescues cardiomyocytes and mitochondria from TKI exposure

## INTRODUCTION

Drug-induced cardiotoxicity is one of the major causes of cardiac diseases, and the underlying mechanisms include mitochondrial dysfunction, altered expression of cardiac genes, and oxidative stress ^1^. Drug exposure during cardiac development increases the risks of developmental cardiotoxicity and congenital heart defect (CHD), which is the most common birth defect affecting nearly 1% of newborns every year ^2,3^. Drug-induced developmental cardiotoxicity exerts significant impacts on the quality of life and increases health care costs in the U.S. However, unlike CHDs caused by chromosomal abnormalities or gene mutations, the mechanisms underlying abnormalities and defects in cardiac development from non-inherited factors (such as maternal exposure to chemicals) is poorly understood, leading to challenges in prediction and prevention of drug-induced developmental toxicity ^4^.

Human stem cells provide a great opportunity for mechanistic and predictive developmental toxicology studies. Cellular differentiation from stem cells can be used to recapitulate embryonic developmental process ^5,6^. Chemical perturbation during stem cell differentiation allows us to understand the impact of drug toxicity on development as well as the underlying molecular mechanisms. We have previously applied human stem cells towards a better understanding of the transcription regulation driving 13-*cis*-retinoic-acid-induced disruption in mesoderm formation ^7^.

In this study, we performed chemical perturbation with various drugs during cardiac differentiation from H1 human embryonic stem cells (H1-ESCs). These drugs were classified as pregnancy category C, D and X by the FDA’s previous version of the Pregnancy and Lactation Labeling Rule (PLLR), meaning that they have been determined to pose a high risk of birth defects in humans. The drugs selected for this study have been reported to exert high risk of CHD, including tyrosine kinase inhibitors (TKIs) for cancer chemotherapy (imatinib, sunitinib and vandetanib; category D) ^8-10^, immunosuppressants (tacrolimus [category C] and mycophenolate [category D]) ^11,12^, and thalidomide (known as teratogen [category X]) ^13,14^. Unlike category X drugs which are not allowed for use in pregnant women, category C and D drugs may be used during pregnancy if clinical benefits outweigh their risks. For instance, anti-cancer drugs (*e.g.*, TKIs) are capable of crossing the placental barrier and posing high risks of CHDs ^9,15^; although cases of pregnant women with cancer are uncommon, with incidences of approximately 1-2/1000 ^16,17^. Single cases of related CHDs from maternal exposure to cancer drugs have been documented in clinical reports ^8,10,18,19^.

Transcription regulatory mechanisms during cardiomyocyte differentiation upon drug exposure were explored in the present study, by integrative network analysis of genome-wide transcriptomics and chromatin accessibility. We discovered a novel a crosstalk mechanism between transcription factor GATA4 and mitochondrial function and biogenesis upon TKI exposure. Results from this study will help us to fill knowledge gaps in our understanding of the mechanisms of cardiac and metabolic dysfunctions due to drug exposure, and will benefit the predictive toxicology for prevention of drug-induced toxicity for human health in the future.

## RESULTS

### Developmental exposure to sublethal level of TKIs induces disarrangement of sarcomere and alteration in Ca^2+^-handling of cardiomyocytes

We utilized a well-established protocol ^20^ for cardiomyocyte (CM) differentiation, which consistently generates high-purity beating CMs derived from human stem cells. In order to optimize the viability of differentiated CMs for the following function analysis, we first determined the no-observed-adverse-effect-levels (NOAELs) of these drugs for CM differentiation by dose-responsive experiments (Supplemental Figure 1A). The NOAEL for TKI drugs (imatinib, sunitinib and vandetanib) was 250 nM, and this is lower than their concentrations in blood based on the clinical data ^15,21-25^. In addition, The NOAELs for tacrolimus and mycophenolate were 50 nM, and the NOAEL for thalidomide was 100 nM. H1-hESCs were then exposed to each drug at the corresponding NOAEL throughout the CM differentiation, and 0.1% of DMSO was used as a vehicle control (Supplemental Figure 1A).

Drug exposures were conducted using two experimental strategies, including: Exposure I, a brief early exposure design, in which the TKI drugs were added from day 0 to day 6 (hereafter referred to as the cardiac progenitor stage ^7^); Exposure II, and a chronic exposure strategy, in which cells were exposed to TKI drugs throughout the entire differentiation process until CMs were collected for analyses (Figure1A). No significant differences in efficiency of CM differentiation were observed between control and drug-treated groups (Figure1B and Supplemental Figure 1B). However, in Exposure II, imatinib and sunitinib (250nM) induced obvious disarrangements of myofilaments of differentiated CMs, whereas 250nM of vandetanib induced only modest adverse effects on sarcomere structure (Figure1C). To evaluate the impacts of drug-exposure on differentiated CMs, spontaneous Ca^2+^ transients were analyzed in the differentiated CMs after day 25. We observed TKI-induced abnormal Ca^2+^-handling in both Exposure I and II, including decreases in beating rates and amplitude, increases in time to peak, TD90, TD50, and impaired calcium recycling (*i.e*., increased decay tau) (Figure 1D-1G). Both Exposure I and II of TKIs exhibited similar effects on Ca^2+^ handling of differentiated CMs.

**Figure 1.**
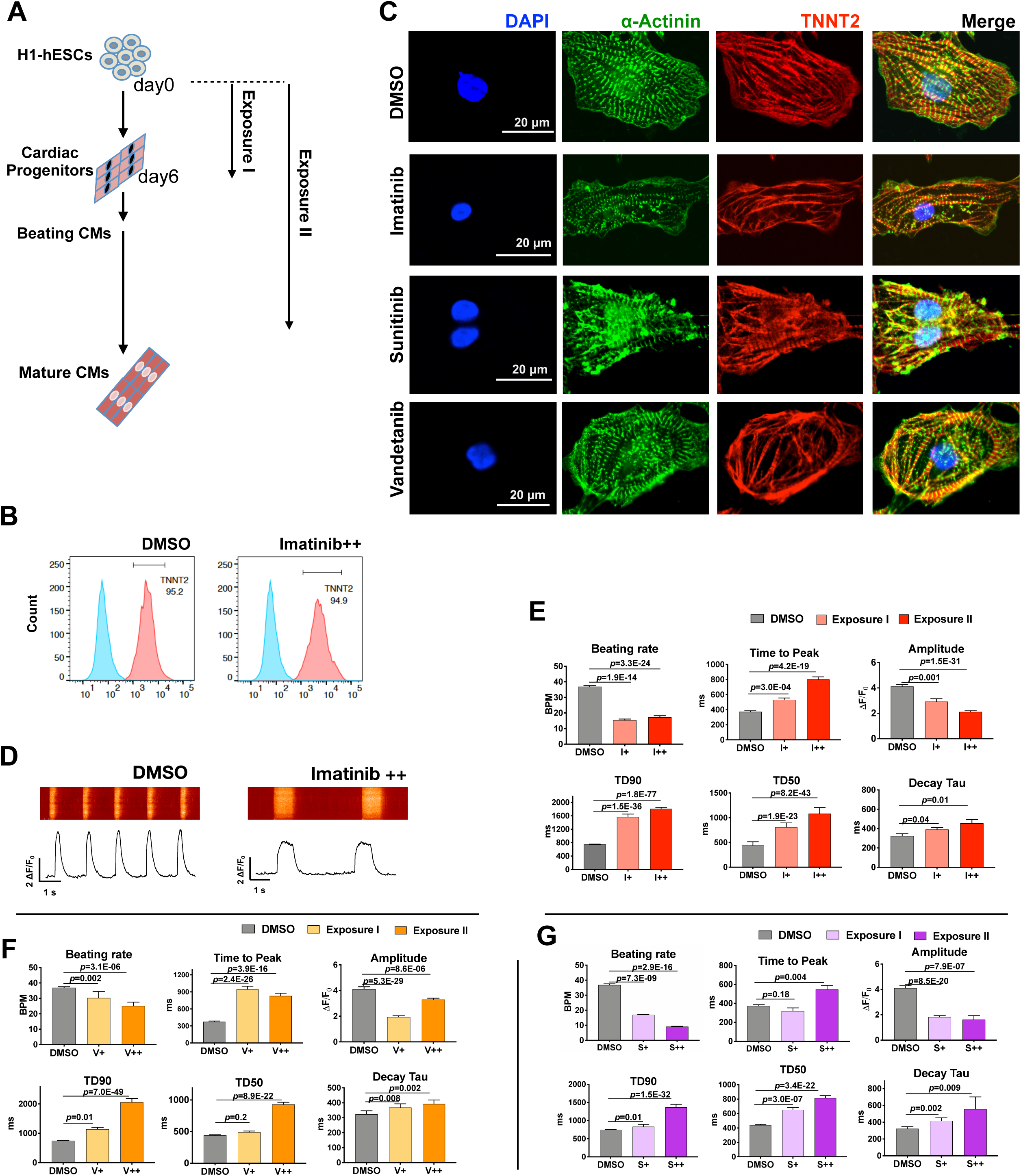
Developmental exposure to sublethal level of TKIs induced disarrangement of sarcomere and alteration in Ca^2+^-handling of cardiomyocytes. **(A)** Experimental design of drug exposure strategies. H1-ESCs were differentiated to cardiomyocytes upon chronic exposure to sublethal of drugs following two exposure designs: Exposure I, the TKI drugs were added from day 0 to day 6 (cardiac progenitor); Exposure II, the cells were exposure to TKI drugs till the differentiated CMs were collected for analyses. **(B)** Analysis of the efficiency of CM differentiation with flow-cytometry of TNNT2-positive cell populations, no significant difference was observed between control (0.1% DMSO) and imatinib groups. More results of other groups are shown in Supplemental Figure 1B. **(C)** Immunostaining of differentiated cardiomyocytes with α-actinin (green), TNNT2 (red), and DAPI (blue). Developmental exposure to TKIs caused disorganization of sarcomere structures of cardiomyocytes from Exposure II. **(D)** Representative Ca^2+^-handling recording from control and imatinib-treated cardiomyocytes. **(E-G)** Bar charts represent developmental exposure to TKIs caused significant alternations of Ca^2+^-handling properties, including decreases in beating rate and amplitude, and increased in time to peak, TD90/50, and decay tau. The *p* values were calculated by nonparametric T-test between control and TKI-treated groups. I, imatinib; V, vandetanib; S, sutininib; +, Exposure I; ++, Exposure II.

To determine whether these toxic effects were specific to anti-cancer TKIs, we also evaluated other drugs associated with maternal-fetal toxicity. We found that NOEALs of tacrolimus, mycophenolic acid, and thalidomide caused dysfunction in Ca^2+^-handling (Supplemental Figure 2A) in differentiated CMs; while no significant morphological change was observed. These results demonstrate that, at NOAELs, developmental exposure to TKI drugs caused more severe effects during CM differentiation than non-TKI drugs used in this study.

### Transcriptomic analysis revealed correlation between the transcriptional regulatory network and mitochondrial function

In order to elucidate the changes in gene expression during CM differentiation due to developmental drug exposure, we performed genome-wide transcriptomic analysis of differentiated CMs (day 20) using RNA-sequencing (RNA-seq) experiments. Twelve modules (*i.e.*, networks) were identified by weighted correlation network analysis (WGCNA) (Figure 2A), and the full lists of genes and enriched GO terms are shown in Supplemental Tables 1 and 2. Clustering analysis of the transcriptomic profiles revealed that TKI drug-treated CMs exhibited high concordance between Exposure I and II (except Exposure I of imatinib) (Figure 2B). We observed that gene expression within a module 1 exhibited down-regulation patterns in TKI-treated groups compared to other groups (*i.e*., immunosuppressant drugs, thalidomide, and 0.1% DMSO), and the representative enriched-GO terms of this module are related to “heart development” and “mitochondrial respiratory chain complex I assembly” (Figure 2C). Genes known to regulate cardiac development were assigned to this module, such as *GATA4, MEF2A, TBX20*, and *HAND2*. In addition, genes involved in oxidative phosphorylation (OXPHOS) (such as *NDUFV3 and NDUFA12*) and glycolysis (such as *HK1 and ALDOA*) were also assigned to this module (Figure 2D). Whereas exposure to TKIs generally led to down-regulated gene expressions in this module, exposure to immunosuppressant and thalidomide drugs had the opposite effect (Figure 2E), suggesting that TKIs induced adverse effects on both cardiac functions and metabolisms *via* dysregulation of transcription factors (TFs) (such as GATA4) and metabolic genes, which exhibited some correlations during CM differentiation.

**Figure 2.**
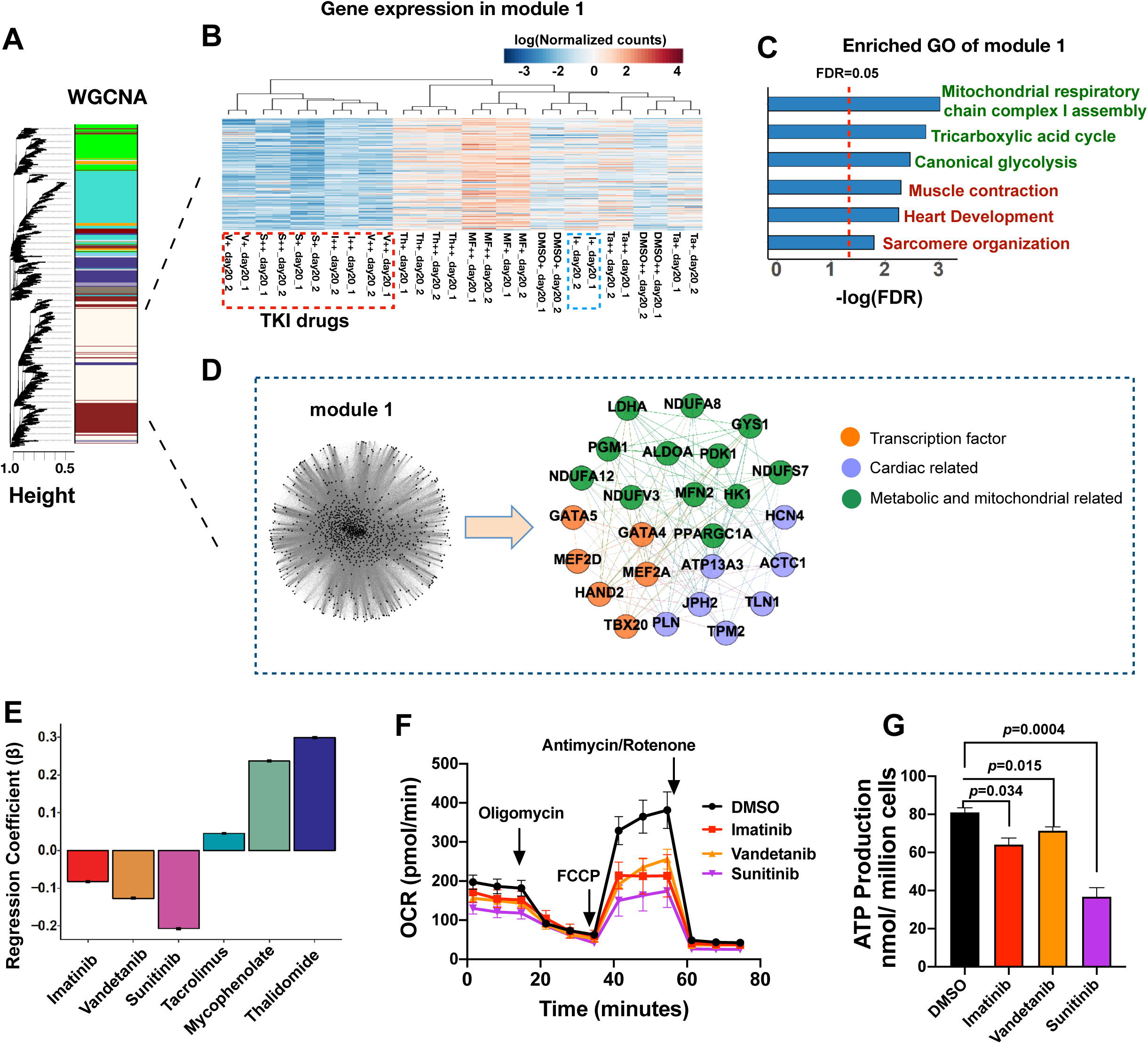
Transcriptomic analysis revealed correlation between transcriptional regulatory network and mitochondrial function. **(A)** Weighted gene co-expression network analysis (WGCNA) revealed modules based on transcriptomic profiles from differentiated CMs (day20) in control and drug-treated conditions. **(B)** The heatmap represents the gene expression pattern of the module 1, using the normalized counts by trimmed mean of M-values (TMM) normalization method. I, imatinib; V, vandetanib; S, sutininib; Th, thalidomide; Ta, tacrolimus; and MF, mycophenolate. Clustering analysis of the transcriptomic profile revealed that TKI drug-treated cardiomyocytes exhibited high concordance between Exposure I (+) and Exposure II (++) (except Exposure I for imatinib) **(C)** Statistically enriched (FDR<0.05, red line indicated) gene ontology (GO) terms of genes in the module 1 (FDR<0.05, red line indicated). **(D)** Representative nodes from module 1, including transcription factor controlling heart development, cardiac genes, and metabolic and mitochondrial related genes. **(E)** Bar chart shows the regression coefficients of different drugs when regressing the mean gene expression of module 1 on the drug usage. The data are normalized to DMSO. **(F)** Evaluation of mitochondrial oxygen consumption rate (OCR) in cardiomyocyte upon exposure to TKIs during differentiation. The data are normalized by cell numbers. **(G)** ATP production was significantly inhibited in differentiated cardiomyocytes by developmental exposure to TKIs. Data are reported as means ± standard error of the mean (SEM). The corresponding *p* values were calculated by nonparametric T-test between control and TKI-treated groups.

Differentiated CMs mainly use OXPHOS to support their large ATP demands ^26-28^; therefore, we examined mitochondrial respiratory activity in TKI-treated CMs. We found that chronic exposure to TKIs decreased the levels of both basal respiration and maximal respiration in differentiated CMs (Figure 2F). The ATP production was also decreased in TKI-treated cells, and sunitinib-treated cells exhibited the lowest ATP level compared to other groups (Figure 2G). By transcriptomic profiling of the genes involved in Ca^2+^-handling, ion channels, and β-adrenergic signaling, we showed that TKIs caused down-regulation of *RYR2, CAMK2D*, and *MYH7*, which are closely related to the regulation of CM contraction (Supplemental Figure 2B). These results suggested that developmental exposure to TKIs dysregulated expression of genes involved in cardiac differentiation, contractile functions, and metabolism; impaired mitochondrial respiration and ATP productions; and ultimately weakened contraction of CMs.

### Motifs enrichment analysis reveals features of transcription-factor occupancy in TKI-treated cardiomyocytes

Many of the genes disrupted by TKI exposure encode key transcription factors important for CM differentiation (Figure 2A). We then leveraged ATAC-seq to identify chromatin accessibility that correspond to altered TF occupancy ^29,30^. Specifically, we measured TF-motif enrichments within differential ATAC-seq peaks between control and TKI-treatment groups, and identified several motifs that are significantly enriched in regions where chromatin accessibility was lost in the TKI-treated groups compared to control (Figure 3A). These enriched motifs correspond to several TF families, that are known to play important roles in CM differentiation (such as GATAs, TEADs, and MEF2 family members) (Figures 3B and Supplemental Figure 3A). The control group exhibited higher densities of motifs for GATA4 and TEAD1/3 (Figure 3C) and stronger ATAC-seq signals (accessibilities) around the summit of the motifs for GATA4 and TEAD1/3, compared to that of TKI-treated cells (Figures 3D-3F; Supplemental Figure 3B). The sunitinib-treated cells showed the most significant loss of chromatin accessibility surrounding the TF motifs, suggesting that sunitinib exposure may have the strongest adverse effects on DNA-binding of these TFs during CM differentiation. Altogether, these results suggest that developmental exposure to TKIs leads to significant changes in the binding patterns of critical developmental TFs, leading to transcriptional dysregulation that may ultimately underlie the toxicity of drug exposure during CM differentiation.

**Figure 3.**
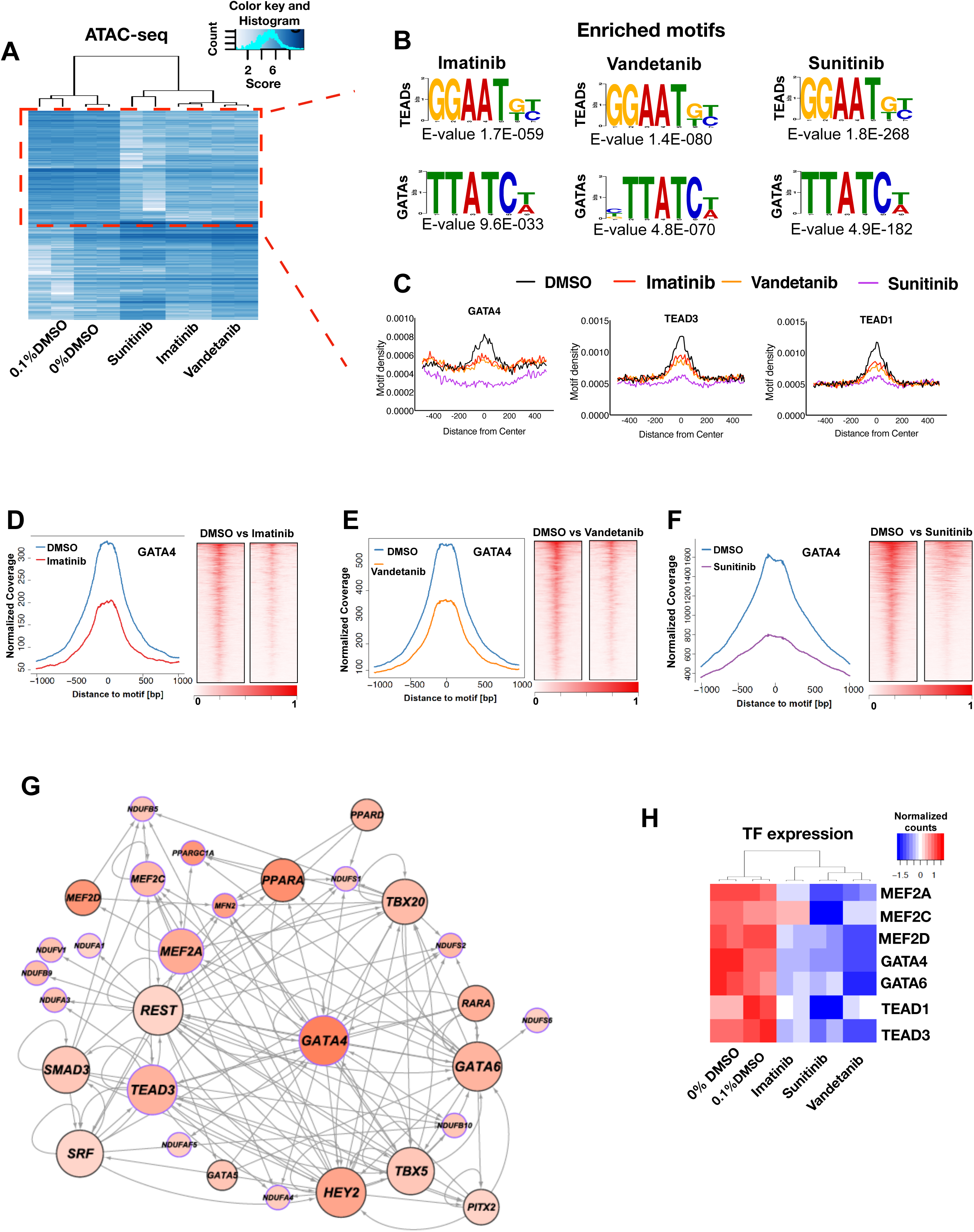
Integrative network analysis revealed strong interactions between transcription factors and mitochondrial genes. **(A)** Genome-wide landscape of chromatin accessibility in differentiated cardiomyocytes between DMSO (0% and 0.1%), and TKIs. The red box indicates the down-regulated chromatin accessibility in the TKI-treated cardiomyocytes. High correlation between 0% DMSO and 1% DMSO indicated that DMSO did not cause aberrations in chromatin accessibility. **(B)** Enriched *de novo* motifs (E value <0.001) were discovered using MEME-chip based on down-regulated chromatin accessibility in TKI-treated cells. **(C)** Motif density plots of representative transcription factors. **(D-F)** ATAC-seq peak signals (+/- 1000 bps) arounds submits of the motifs for GATA4 in TKI-treated cells compared with DMSO-treated cells. Binding intensities are shown as sequencing depth-normalized tag count. These figures are partial data of Supplemental Figure 3B. **(G)** Visualization of most representative genes from the integrative transcriptional regulatory networks. Both RNA-seq and ATAC-seq data were integrated to uncover the transcriptional regulatory networks with the paired expression and chromatin accessibility (PECA) model^31^. The size of the node represents the out-degree, which is the number of target gene. The color of the node represents the expression fold-change. **(H)** Gene expression of selective TFs shown in the Figure 3G.

### Integrative genome-wide transcriptomic and open-chromatin analyses revealed GATA4 as an important regulator for TKI exposure

We further integrated both chromatin landscapes and gene expression data to explore transcriptional regulatory networks (Figure 3G), using paired expression and chromatin accessibility (PECA) model, which can elucidate the effect of TFs bound to the activated context-specific *cis*-regulatory elements on the transcription of target genes ^31,32^. We found that genes within the module 1 from the WGCNA, which are involved in mitochondrial biogenesis and function, are predicted to be regulated by transcription factors known to control heart development, such as GATA4, MEF2A/C and TEAD1/3 (Figure 3G). These genes include peroxisome proliferator-activated receptor alpha (*PPARA*) and peroxisome proliferator-activated receptor gamma coactivator 1-alpha (*PPARGC1A*, encoding PGC-1α, which is a key regulator for mitochondrial biogenesis) ^33-35^, complex I (*i.e*., *NDUF* gene family), and cardiac development (such as *TBX5*) ^36^ (Figure 3G). Importantly, Expression levels of these TFs were down-regulated in differentiated CMs upon exposure to TKIs (Figure 3H). In particular, *GATA4* exhibits high degree (*i.e*., more targeted genes likely regulated by GATA4) within the network, indicating important roles of GATA4 in TKI-induced transcriptional dysregulation.

Mitochondrial biogenesis plays an important role during heart development, and heart development is highly associated with mitochondrial dynamics, biogenesis, and oxidative metabolism^28,37-39^. A previous study showed that genomic GATA4-occupied regions in mouse fetus (E11.5) are associated genes related to “mitochondrial organization”, such as *Ppargc1a* ^40^, suggesting that GATA4 may be one important regulatory factor for mitochondrial complex assembly during prenatal development. In our study, the WGCNA analysis, the motif enriched analysis, and the PECA analysis strongly suggest that *GATA4* is likely to regulate genes involved mitochondrial complexes and biogenesis. Thus, we hypothesize that TKIs induce mitochondrial dysfunction during CM differentiation *via* inhibition of GATA4-mediated networks, ultimately leading to dysfunctions in differentiated CMs. This led us to investigate the role of GATA4 in CM differentiation upon exposure to TKI, and to examine whether CMs and their mitochondrial function can be restored through overexpression of *GATA4.*

### Gain-of-function by GATA4 overexpression restores cardiomyocyte functions in the presence of TKI exposure

In order to better understand the role of GATA4 in TKIs induced cardiotoxicity during differentiation, we created a lentiviral delivery-based dCas9/CRISPR-activation (dCas9/CRISPRa) system based on previous studies ^41,42^, which allows us to find out whether enhance of GATA4 expression will ameliorates TKI-induced toxicity in differentiated CMs. The GATA4 overexpression *via* this dCas9/CRISPRa system is doxycycline-inducible (Figure 4A and Supplemental Figure 4A). Because our RNA-seq data showed that GATA4 expression was down-regulated after day 6 upon exposure to TKI drugs (Supplemental Figure 4B), 2 μg/ml doxycycline was added daily to induce the expression GATA4 after day 6 (Figure 4B-4D). In spontaneous calcium transient analysis, we found the impaired Ca^2+^-handling signaling of TKI-treated CMs was restored by GATA4 induction, including increased beating rate and amplitude, faster calcium handling as the time to peak, TD90, TD50, and decay tau were all decreased (Figure 5A-5D). In addition, we also observed the disorganization of sarcomeres in TKI treated CMs were reinstated by GATA4 expression (Figure 5E). These results suggest that TKI-induced toxicity in CMs can be reversibly recovered by enhancing GATA4 expression.

**Figure 4.**
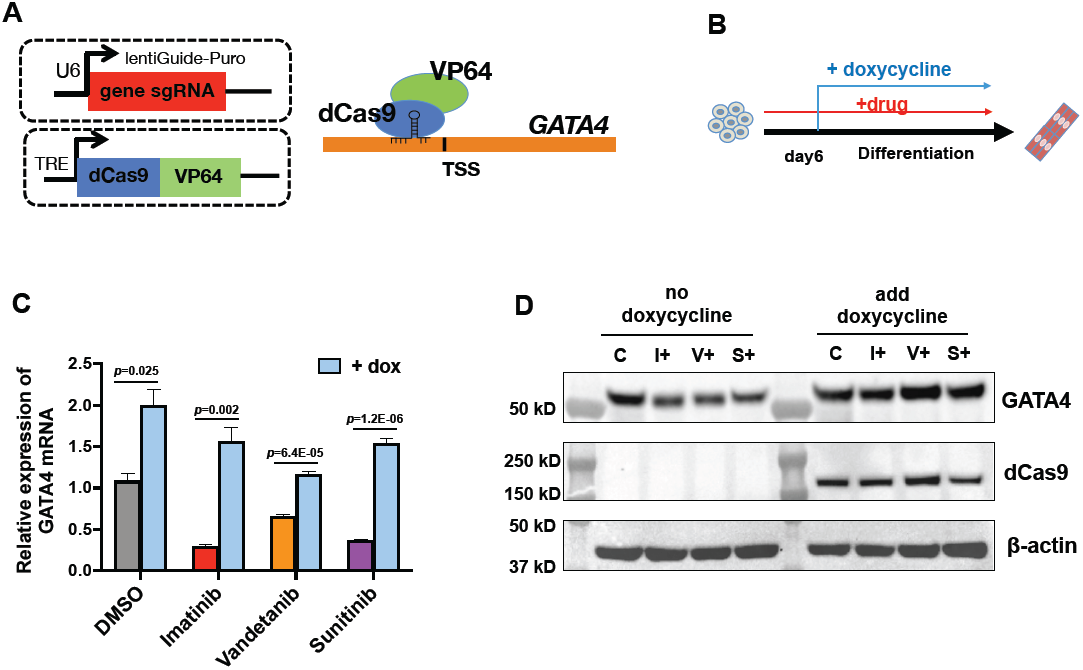
Design of CRISPR-activation system for GATA4. **(A)** Schematic of the constitutive TRE-regulated dCas9-VP64 and GATA4 sgRNA constructs. **(B)** GATA4 was overexpressed by adding doxycycline (dox) from day 6 in presence of TKIs during cardiac differentiation of H1-hESCs. **(C)** Quantitative gene expression analysis of tetracycline response element (TRE)-regulated dCas9-VP64 cells transduced with *GATA4* sgRNAs. Data are reported as means ± standard error of the mean (SEM). The *p* values were calculated between cells with and without adding dox. **(D)** Expression of GATA4 protein on in differentiated CMs using Western blot analysis.

**Figure 5.**
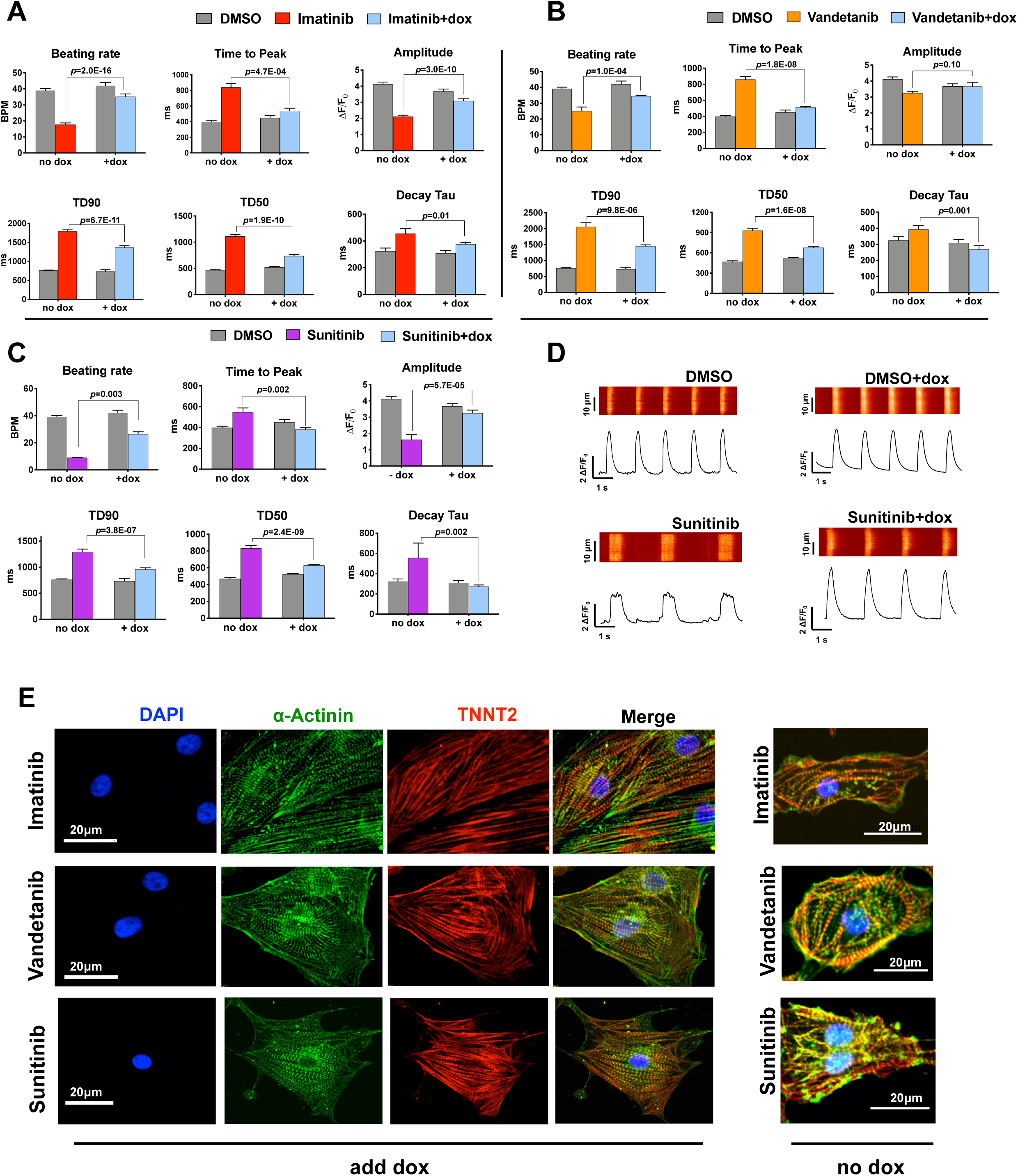
Overexpression of GATA4 restored Ca^2+^-handing and reinstated sarcomere structures in differentiated cardiomyocytes upon developmental exposure to TKIs. **(A-C)** Bar charts represent alternations of Ca^2+^-handling properties in TKI-treated cardiomyocytes after induction of *GATA4* expression, including increases in beating rate and amplitude, and decreased in time to peak, TD90/50, and decay tau. Data are reported as means ± standard error of the mean (SEM). The *p* values were calculated between TKI-treated cells with and without adding doxycycline (dox). **(D)** Representative Ca^2+^-handling traces from DMSO and sunitinib-treated cardiomyocytes with and without GATA4 induction. **(E)** Immunostaining of differentiated cardiomyocytes with α-actinin (green), TNNT2 (red), and DAPI (blue). Disarrangement of sarcomere structures in TKI-treated cardiomyocytes were reinstated after induction of *GATA4* expression.

### Overexpression of GATA4 restored mitochondrial function in differentiated cardiomyocytes upon TKI exposure

Integrative network analysis showed that GATA4 is involved in the regulatory networks related to cardiac functions, metabolic process, and mitochondrial function, and TKI-induced impairment of mitochondrial respiration. We therefore investigated whether TKI-impaired mitochondrial respiration of CMs can also be restored by enhancing GATA4 expression.

As expected, we observed overexpression of GATA4 during CM differentiation improved basal and maximal respiration in differentiated CMs in the presence of chronic exposure to the TKIs (Figure 6A-6C). This enhancement of mitochondrial respiration by overexpression GATA4 was also observed in the control group in the absence of drug exposure (Figure 6D). We also found that overexpression of GATA4 promoted ATP production (Figure 6E) and mitochondrial membrane potential (Figure 6F) in the CMs upon developmental exposure to TKIs, and the most significant improvement was observed in sunitinib-treated group. Moreover, chronic TKI exposure decreased branch size of the mitochondria in differentiated CMs compared to control, which was reversed by induction of GATA4 (Figure 6G-6O).

**Figure 6.**
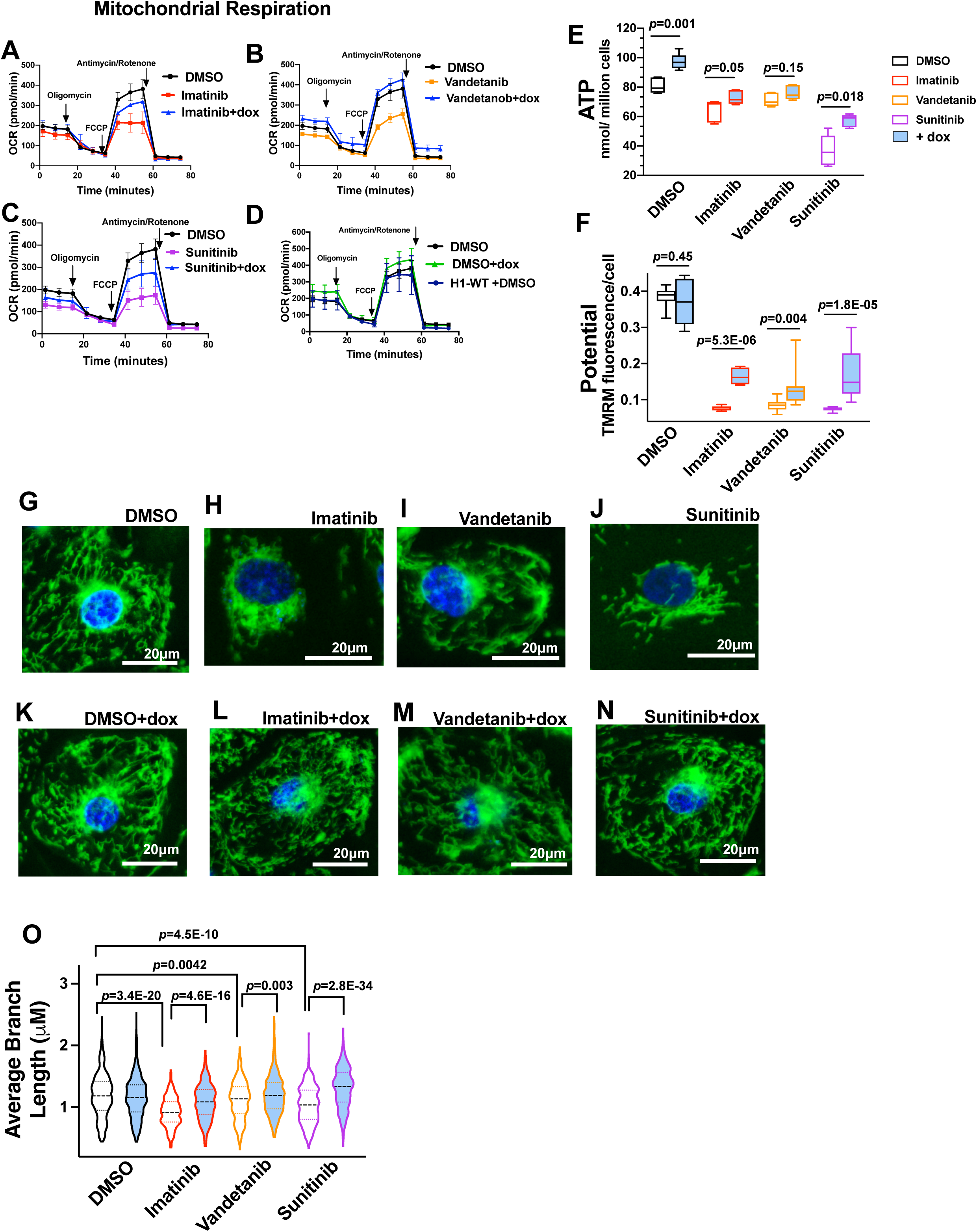
Overexpression of GATA4 restored mitochondrial function and change mitochondrial morphology in differentiated cardiomyocytes. **(A-C)** Overexpression of *GATA4* enhanced mitochondrial oxygen consumption rate (OCR) in TKI-treated cardiomyocytes. The data were normalized by cell numbers. **(D)** Enhanced mitochondrial respiration was also shown in control group (0.1% DMSO) by induction of *GATA4* expression. H1-hESCs (wide type, H1-WT) and H1-hESCs carrying CRISPR-a for GATA4 showed similar OCR without adding doxycycline. **(E-F)** Overexpression of *GATA4* increased ATP production (E) and mitochondrial membrane potential (F) in TKI-treated cardiomyocytes. The *p* values were calculated between TKI-treated cells with and without adding doxycycline (dox). **(G-J)** Developmental exposure to TKIs caused smaller branch sizes in differentiated cardiomyocytes (H-J) compared to control group (G). **(K-N)** Overexpression of GATA4 increased the branch sizes in differentiated cardiomyocytes. Mitochondria in live cells were stained with MitoTracker dye (green). Nucleus were stained with DAPI (Blue). **(O)** Quantitative analysis of mitochondrial branch sizes using the Fiji software with a published method ^96^.

We next examined dynamics of mitochondrial DNA (mtDNA) levels during CM differentiation, and found that mtDNA numbers increase from cardiac progenitor stage (day 6) towards to the differentiating CM stages (Supplemental Figure 5A), suggesting similarity in the increases of cardiac mitochondria during both *in-vivo* and *in-vitro* cardiac differentiation ^38,43^. Chronic TKI exposure significantly relatively lowered mtDNA copy numbers compared to control cells after day 6 of differentiation compared to the control group (Supplemental Figure 5B-5C), whereas overexpression of GATA4 increased the mtDNA copy numbers (Supplemental Figure 5D), suggesting that overexpression of GATA4 promotes mitochondrial biogenesis.

We explored the genome-wide GATA4-binding sites using ChIP-seq experiments, and we found a global loss of GATA4-binding upon sunitinib exposure compared to that in control group (Figure 7A). Genes regulated by GATA4 are related to cardiac structure (“actin filament binding” and “PDZ domain binding”), homeostasis of ion channels (“ion channel binding”), and signaling pathways (“SMAD binding”). By contrast, sunitinib exposure attenuated the enrichment of relevant GO terms in drug-treated CMs, consistent with loss of GATA4-binding within these regions (Figure 7B), demonstrating sunitinib induced adverse effects on cardiac dysfunction and sarcomere structure *via* inhibition of transcriptional activity of GATA4.

**Figure 7.**
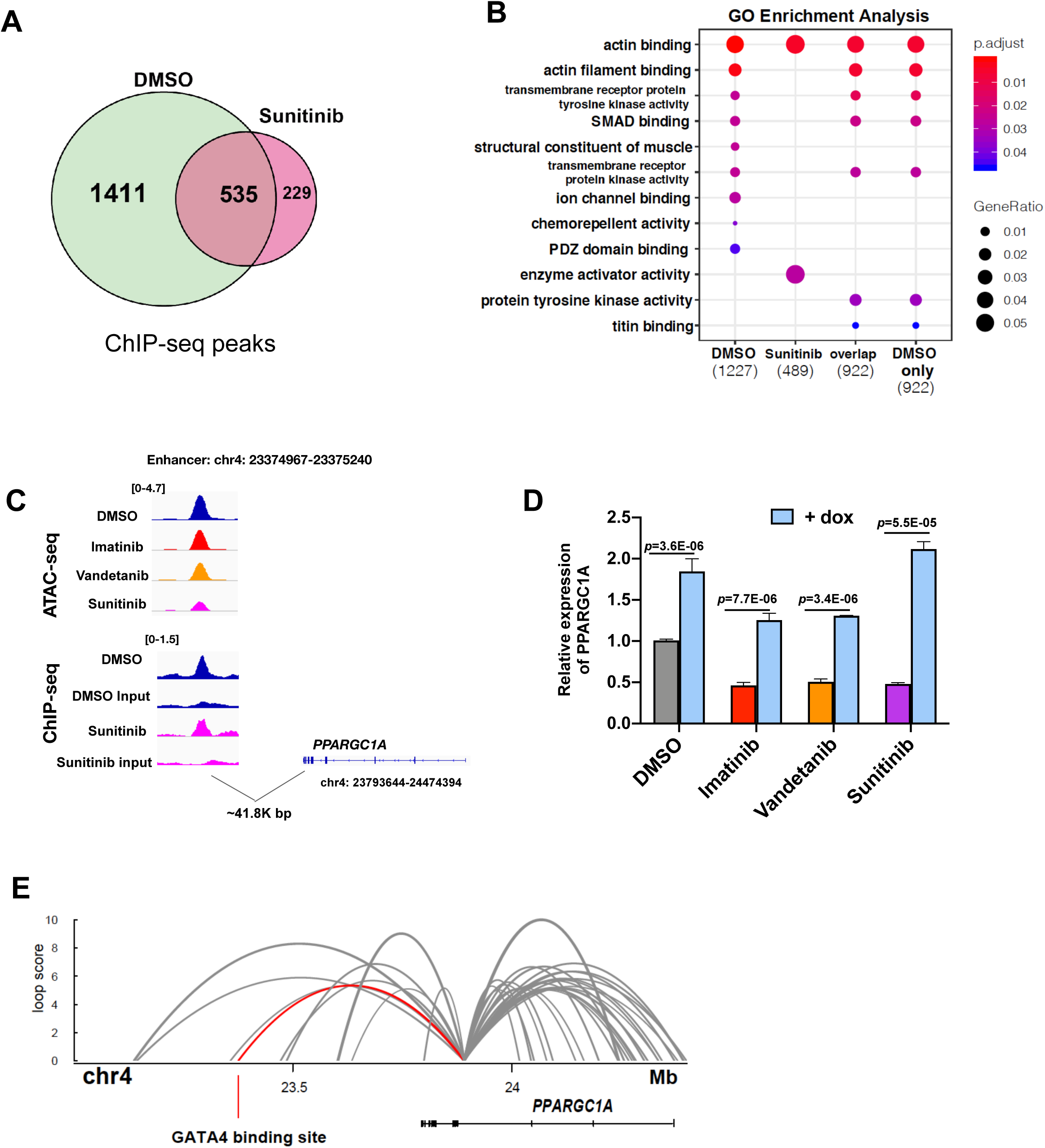
GATA4 occupancy analysis by ChIP-seq experiments. **(A)** Venn diagram represents numbers of genomic DNA-binding locations of GATA4 between DMSO and sunitinib. (**B)** GO enrichment analysis of genes close to +/- 3000 bps of GATA4-binding sites corresponding to the Venn diagram (A). **(C)** Both ATAC-seq and ChIP-seq of GATA4 data showed higher signals at enhancer region of PPARGC1A. **(D)** Quantitative gene expression analysis of PPARGC1A. Data are reported as means ± standard error of the mean (SEM). The dox represents doxycycline. (**E**) Visualization of long-range *PPARGC1A* promoter interactions in iPSC-derived cardiomyocytes confirms direct interaction between GATA4-binding site and PPARGC1A anchor region. Red loop specifies individual long-range interaction called between the GATA4 binding site coordinates and the associated anchor used to capture PPARGC1A promoter interactions in cardiomyocytes ^45^.

Previous studies have showed that GATA4-binding DNA regions function as enhancer regulatory elements for *Ppargc1a*, and overexpression of GATA4 can increase *Ppargc1a* promoter activity ^40,44^. In the present study, specific analysis of GATA4 occupancy near *PPARGC1A* identifies a GATA4 binding site approximately 42-Kb upstream of *PPARGC1A* (Figure 7C and Supplemental table 3). ChIP-seq experiments carried out in drug-treated CMs identifies a loss of GATA4 occupancy at this putative *PPARGC1A* enhancer in sunitinib-treated CMs compared with control (Figure 7C). Both mRNA and protein expression levels of PGC-1α were downregulated in differentiate CMs upon TKI exposure and, conversely, upregulated by overexpression of GATA4 (Figure 7D and Supplemental Figure 5E). Moreover, further integration of the current GATA4 ChIP-seq data with promoter-capture Hi-C data previously reported in analogous day 20 CMs ^45^, we confirmed a direct interaction between the GATA4-binding site with the *PPARGC1A* promoter anchor in CMs, consistent with long-range enhancer-promoter regulation through DNA looping (Figure 7E). Given the important roles of PGC-1α in regulating mitochondrial biogenesis and functions, altogether, this demonstrates that GATA4 binds to a *PPARGC1A*-enhancer and directly influences the expression of PGC-1α, thereby controlling mitochondrial biogenesis and function at early stages.

### Maternal exposure to sublethal level of sunitinib induced developmental defects in mouse fetus heart

To evaluate the *in-vivo* toxicity of sunitinib during cardiac development, we investigated the effects of maternal sunitinib exposure in mouse fetus. Concentration of sunitinib used for daily intraperitoneal injection in female mouse was 3mg/kg/day, which is proportional to the concentration used for human stem cells in this study. Mice embryos were exposed to sunitinib from prenatal stages (E0) towards neonatal stage (P0) (Supplemental Figure 6A). We observed that maternal sunitinib exposure caused embryonic death (2 cases, Supplemental Figure 6B) and moderate CHD-like histopathologic morphology (*i.e*., thinner myocardium and ventricular septal defect) in some of the surviving pups (around 10%) (Supplemental Figure 6C-6E). We also observed high level of bioaccumulation of sunitinib in the blood of both female mice and newborns (P0), and level of sunitinib in P0 newborns was 80% of that in female (Supplemental Figure 6F). These results suggest that TKIs can accumulated in embryos by maternal exposure and exert developmental toxicities during fetal development.

## DISCUSSION

### Human stem cells as a cell-based platform to study developmental cardiotoxicity

The present study provides a new paradigm of using human stem cells to model developmental exposure, so as to better understand the mechanisms underlying drug-induced developmental cardiotoxicity. It has been reported that exposure to various chemicals during pregnancy increases the risks for CHDs, and many of these chemicals are able to cross the placental barrier ^46^, such as heavy metals (e.g., lead ^47^ and cadmium ^48^), pesticides ^49,50^, arsenic ^48^. In addition, a number of medications are known to have high risks for congenital defects, such as thalidomide ^13,14^ and retinoic acids ^51-53^. Due to ethnical restrictions curtailing the use of human embryonic tissues in scientific researches, human stem cells have provided us an alternative model to understand developmental processes in humans ^5,6^.

### TKIs and developmental cardiotoxicity

TKI drugs have been used for cancer chemotherapy. However, they are known to cause adverse effects in cardiovascular system in adults (*i.e.*, off-target toxicity), including hypertension, left ventricular systolic dysfunction, QT prolongation, and heart failure ^54-57^. As TKIs can cross the placenta to expose the fetus during development ^15^, TKI-induced developmental cardiotoxicity has also been reported in human ^8-10,18^ and animals ^58^. On the other hand, mutation of several targets for TKIs are associated with CHD, such as ABL Proto-Oncogene (*ABL1*, targeted by imatinib) ^*59*^, and platelet-derived growth factor receptor alpha (*PDGFRA*, targeted by imatinib and sunitinib) ^60,61^. All these reports strongly suggested that maternal exposure of TKIs present high risks of developmental cardiotoxicity and CHD.

The concentration of 250 nM used in this study was lower than plasma levels of these TKI drugs achieved in clinical use ^15,21-25^. Our results from modeling stem cells provides direct evidences of cardiotoxicity derived from exposure at this concentration, suggesting that sublethal level of TKIs (especially sunitinib) still exert mild toxicity in developing heart and cause dysfunctions (*su*ch as congenital arrhythmia), even though cardiac development still successfully proceeds to four-chamber heart structures in the fetus. The bioaccumulation and cardiotoxicity from maternal sunitinib exposure was evident in the *in-vivo* mouse study, and the potential long-term adverse effects of sunitinib in offspring need to be characterized in future studies.

### GATA4 as a key regulator in cardiotoxicity from exposure to anticancer drugs

Previous studies showed that overexpression of GATA4 in early developmental stages promote CM differentiation ^62,63^. Although overexpression of GATA4 in the neonatal rat causes a mild form of cardiac hypertrophy at six months of age ^64^, GATA4 acts as important regulator for cardiac angiogenesis and myocardial regeneration in neonatal mice ^65,66^. These studies suggest the multiple roles of GATA4 in both early and postnatal cardiac developmental stages. Moreover, GATA4 has been shown to play protective roles in cardiotoxicity from exposure to anthracycline anticancer agents (*e.g*., doxorubicin), and overexpression of GATA4 can reduce doxorubicin-induced apoptosis and increase the survival of CMs ^67,68^. Regarding effects of TKIs on GATA4, a single study by Maharsy *et al*. ^69^ showed that down-regulation of GATA4 in heart is associated with dietary imatinib exposure in mice. Whether GATA4 could exert protective roles in TKI-treated CMs was unknown.

In our study, both gene and protein expression, as well as DNA-binding of GATA4, were attenuated upon exposure to TKIs, demonstrating that GATA4 was targeted by TKIs, similar to doxorubicin, consequently leading to dysregulation of GATA4-mediated transcription networks and cardiac dysfunction in differentiated CMs. In addition, we also observed variation in inhibition of TFs among different TKI drugs. For instance, sunitinib exerted the most significant adverse effects on genomic binding of GATA4, compared to that of imatinib and vandetanib; while vandetanib exerted more significant effects on MEF2s (Supplemental Figure 3A). Giving the obvious toxic effects (*e.g*., sarcomeric disarrangement, mitochondrial size, and ATP production) on sunitinib-treated group, this demonstrates that GATA4 acts as important role in protection of sunitinib-induced developmental toxicity.

### Mitochondria, GATA4 and cardiac development

It is known that heart development undergoes metabolic switch from glycolysis to OXPHOS^26^. Mitochondrial biogenesis and maturation occur during prenatal stages of mice when glycolysis is dominant. During cardiac development, increases in both cardiac mitochondrial number and inner folding membranes (*i.e.*, cristae) are shown from E8.5 in mice, and mitochondrial complexes started to assemble at E11.5; from E13.5 to P0 in mice, mitochondria are functionally matured, thus the newborns can generate ATP relied on OXPHOS at postnatal stages 38,39,43,70.

TKI drugs haven been shown to induced mitochondrial dysfunction, such as respiration or membrane potential, in variable cell types ^71-73^, suggesting that disruption of mitochondrial biogenesis and functions upon exposure to TKI can cause developmental problems in heart development. A previous study by Kasahara *et al.* ^*74*^ showed that depletion of mitochondrial fusion proteins Mitofusin 1 and 2 (MFN1 and MFN2) arrested heart development in mouse embryos and CM differentiation from embryonic stem cells, and this event was accompanied with the decreased GATA4 levels, suggesting a potential relationship between mitochondrial biogenesis and GATA4 during heart development. However, given both GATA4 and mitochondrial biogenesis are well-known to control heart development, the relationship between GATA4 and mitochondria has not been well-documented. Our study showed that GATA4 occupies enhancer regions for *PPARGC1A* in differentiating CMs, corroborating a previous study using the mouse fetus ^40^. Induction of *PPARGC1A* expression *via* GATA4 overexpression can be considered as a potential approach, which is capable of restoring the mitochondrial respiration and morphology from TKI exposure.

Besides GATA4, the TEADs and MEF2s families are also known to regulated cardiac development ^75-77^; and these TFs are reported to be involved in regulating mitochondrial biogenesis *via* activation of PGC1α expression ^78-80^. Previous studies showed that DNA-binding of TEADs and MEF2s co-occupies with that of GATA4 in mouse fetus, indicating interactions among these TFs in the regulation of genes controlling heart development and mitochondrial biogenesis ^40,81,82^. In the present study, the TKI-treated groups exhibited decreased DNA-binding signals of GATA4, TEADs and MEF2s than that in control group, suggesting that TKIs exposure also targeted TEADs or MEF2s in differentiating CMs. Given the importance of these TFs in cardiac differentiation and mitochondria, combinatorial overexpression of these TFs could be considered as a more powerful approach to rescue CMs and mitochondrial function from developmental TKI-exposure.

In sum, we demonstrated cardiotoxicity and mitochondrial dysfunction from developmental exposure to TKI exposure during CM differentiation derived from human stem cells. We demonstrate that TKI-induced toxicity was also observed in mice study. Induction of GATA4 with CRISPRa has restored functions and morphology of differentiated CMs and mitochondria. The present study established an integrated stem cell-omics platform to investigate important mechanisms underlying developmental heart defects from chemical exposure. The novel crosstalk mechanism between GATA4 and mitochondria identified from this study will help us to develop therapeutic solutions to minimize toxicity from maternal drug exposure during early cardiac development.

## EXPERIMENTAL PROCEDURES

### Chemicals

Imatinib Mesylate (S1026, Selleck Chemicals, TX), Sunitinib (RS046, BIOTANG Inc, MA), vandetanib (RS051, BIOTANG Inc), tacrolimus (RS047, BIOTANG Inc), Mycophenolate Mofetil (S1501, Selleck Chemicals), and thalidomide (ICN15875380, Fisher scientific) were ordered and dissolved in DMSO as stocks.

### Cell culture, cardiomyocyte differentiation and chemical exposure

The H1-hESCs (RRID: CVCL_9771) were obtained from the Stem Cell Core Facility of Genetics, Stanford University. The pluripotent cells were grown in Matrigel (Corning)-coated 12-well plates in Essential 8 Medium (Thermo Fisher Scientific) at 37°C incubators (5% CO2). Cardiomyocyte differentiation was initiated using a monolayer differentiation chemically defined method ^20^. Drug exposures were conducted using two experimental designs, including: Exposure I, a brief early exposure design, in which the TKI drugs were added from day 0 to day 6 (cardiac progenitor); Exposure II, in which cells were exposed to TKI drugs throughout the entire differentiation process until CMs were collected for analyses (Figure1A). To further increase cardiomyocyte purity, the differentiated cells were subjected to subsequent glucose starvation using non-glucose-supplemented RPMI/B27 medium twice (two days per time) to decrease non-cardiomyocyte cells, since cardiomyocytes are more tolerant to glucose starvation ^83^. Differentiated beating cardiomyocytes were harvested by TrypLE Select 10X (Thermo Fisher Scientific). Regarding imaging and functional analyses, cells were re-plated with RPMI/B27 supplemented with 10% FBS and 10 μM ROCK inhibitor.

### Immunostaining of sarcomere of differentiated cardiomyocytes

For immunostaining imaging of sarcomere structure, the differentiated cardiomyocytes were re-plated in Nunc™Lab-Tek™II glass-bottomed 8-chamber glass slides (Thermo Fisher), and then cells were fixed and permeated in the plate using a Human Cardiomyocyte Immunocytochemistry Kit (Thermo Fisher Scientific). The primary antibodies included: rabbit anti-cardiac troponin T (Abcam, ab45932, RRID: AB_956386) and mouse anti-α-actinin (sarcomeric) (Sigma-Aldrich, A7811, RRID: AB_476766). The secondary antibodies included: goat anti-rabbit IgG, Alexa Fluro 594 (Thermo Fisher Scientific, R37117, RRID: AB_2556545) and goat anti-mouse IgG, Alexa Fluro 488 (Thermo Fisher Scientific, A-11001, RRID: AB_2534069). The images were taken using Leica DMi8 Microsystems and Zeiss LSM710 inverted confocal microscope, and then the images were processed using the Fiji software (RRID: SCR_002285).

### Spontaneous Ca^2^+ transient imaging and measurement in cardiomyocytes

Differentiated beating cardiomyocytes were dissociated by TrypLE Select 10X (Thermo Fisher Scientific) and 50,000 of cells were re-plated in Matrigel (BD Bioscience) pre-coated 8-well LAB-TEK® II cover glass imaging chambers (Thermo Fisher Scientific). Cells were recovered for 3-4 days after seeding until beating normally. Calcium imaging was performed as previously described ^84^. Briefly, cells were loaded with 5 μM Fluo-4 AM in Tyrode’s solution (140 mM NaCl, 1 mM MgCl_2_, 5.4 mM KCl, 1.8 mM CaCl_2_, 10 mM glucose, and 10 mM Hepes pH = 7.4 with NaOH at RT) for 5-10 min at 37°C. After washing with pre-warmed Tyrode’s solution 3 times, cells were immersed in Tyrode’s solution for 5 min prior to imaging. Spontaneous calcium signaling in cardiomyocytes was sampled by confocal microscopy (Carl Zeiss, LSM 510 Meta, Göttingen, Germany) with a 63X oil immersed objective (Plan-Apochromat 63x/1.40 Oil DIC M27). Signaling was captured in line-scanning mode (512 pixels X 1920 lines) at a speed of 3.2 μs/pixel. For analysis of the data, a custom-made script based on IDL (Interactive digital language) was used. Extracellular background signal was subtracted from calcium signals, and the calcium signal was normalized to the intracellular basal line (F0). Transient amplitude was expressed as ΔF/F0. Decay Tau (mS) was calculated by mono exponential curve fitting.

### Flow cytometry and analysis

Differentiated cardiomyocytes (day20, around 50,000 cells) were collected for flow cytometry. The cells were washed with DPBS buffer (Thermo Fisher Scientific), and then were fixed and permeabilized using Cytofix/Cytoperm (BD Biosciences). Afterwards, the cells were labeled with rabbit anti-cardiac Troponin T (*i.e.* cardiac TNNT2) antibody (Abcam, ab45932, RRID: AB_956386) (1:100 dilution in Perm/Wash buffer, BD Biosciences), and then labeled with goat ant-rabbit IgG (Alexa Fluor 488 conjugated, Thermo Fisher Scientific, A11034, RRID: AB_2576217) secondary antibody (1:200 dilution). Ice-cold DPBS (with 1% FBS) was used as the flow cytometry buffer for re-suspending cells. Flow cytometry was performed using a FACSAria II cytometer (BD Biosciences). The data were analyzed using FlowJo software (version 10.1). The events were first gated to filter dead cells and debris. Troponin T-positive cells were defined as cells having a fluorescence density greater than the isotype control.

### RNA-seq and data analysis

Cells at Day 0, day 6, and day 20 were collected for RNA-seq. For each treatment and each time-point, cells from two independent differentiation wells were used as two biological replicates. Total RNA was extracted from the same number of cells among each group using QIAzol lysis reagent (Qiagen), and RNA was then subjected to DNase I digestion and purified using a miRNeasy Mini Kit (Qiagen) according to the manufacturer’s instructions. RNA integrity was checked with a NanoDrop, and only samples with a ratio of 260/280 between 2.0 - 2.1 were subsequently used for ribosome depletion. Purified RNA (2.5 μg) was used for ribosome depletion using a Ribo-Zero™ Gold Kit (Human/Mouse/Rat) (Epicentre Biotechnologies) according to the manufacturer’s instructions. The integrity of ribosome-depleted RNA was assessed using an Agilent RNA 6000 Pico Assay kit on the Agilent 2100 Bioanalyzer (Agilent Technologies). RNA-seq libraries were constructed using ScriptSeq™ v2 RNA-Seq Library Preparation kits (Epicentre Biotechnologies) according to the manufacturer’s instructions. The concentration of the library was measured with a Qubit Fluorometer (Thermo Fisher Scientific) and the size was determined using an Agilent High Sensitivity DNA kit on an Agilent 2100 Bioanalyzer. All RNA-seq libraries were sequenced by HiSeq4000 sequencers (Illumina) with 2 x 101 cycles.

The raw RNA-seq raw data were trimmed to remove the adapter sequences (including GATCGGAAGAGCACACGTCTGA and AGATCGGAAGAGCGTCGTGTAG) with command-line tool cutadapt (1.8.1). Then the trimmed files were aligned with Tophat (version 2.0.9) to GRCh37/hg19 Homo sapiens reference genome. The human gene symbols and their raw counts were calculated using the HTSeq ^85^ (version 0.6.1p1) package in Python with the hg19 *Homo sapiens* gtf file. Differential gene-expression analysis was performed using the edgeR package in R, and the normalization was performed using a trimmed mean of M-values (TMM) method across all samples ^86^. The Gene Ontology (GO) enrichment analysis was performed using on-line tools DAVID (version 6.8) (https://david.ncifcrf.gov/summary.jsp) and the Gene Ontology Resource (http://geneontology.org). The Gene Ontology (GO) enrichment analysis of differentially expressed genes was performed using DAVID (https://david.ncifcrf.gov).

### ATAC-seq and data analysis

The ATAC-seq protocol developed was used for the chromatin accessibility profiling ^30^. Cells at day 6 and day 20 were collected for ATAC-seq. For each treatment and each time-point, cells from two independent differentiation wells were used as two biological replicates. For each sample, 50,000 cells were collected and pelleted by centrifugation for 15 min at 500 g and 4°C. Cell pellets were washed once with ice-cold 1x PBS and then pelleted again by centrifugation at the previous settings. Cell pellets were re-suspended in 25 μl of ice-cold lysis buffer (10 mM Tris-HCl pH 7.4, 10 mM NaCl, 3 mM MgCl_2_, 0.1% Igepal CA-630), and nuclei were pelleted by centrifugation for 30 min at 500g, 4°C. Supernatants were discarded and nuclei were re-suspended in 50 μl reaction buffer (2.5 μl of Tn5 transposase and 25 μl of TD buffer from a Nextera DNA Library Prep Kit from Illumina, and 22.5 μl nuclease-free H_2_O). The reaction was incubated at 37°C for 30 min, and subsequently the reaction mixture was purified using MinElute PCR Purification Kit (Qiagen). The purified transposed DNA was amplified with NEBNext High-Fidelity 2 X PCR Master Mix (New England Biolabs) and custom-designed primers with barcodes.^30^ Gel electrophoresis was used to remove primer dimers from the PCR products with 2% E-Gel EX Agarose Gels (Thermo Fisher Scientific), and then the PCR products were purified using QIAquick PCR Purification Kit (Qiagen). DNA concentration was measured with a Qubit Fluorometer (Thermo Fisher Scientific) and library sizes were determined using Agilent High Sensitivity DNA kit on Agilent 2100 Bioanalyzer. The ATAC-seq libraries were sequenced with a HiSeq 4000 sequencer (Illumina) with 2 X101 cycles, and the sequencing quality control was performed by Stanford Center for Genomics and Personalized Medicine.

The raw data were trimmed to remove the adapter sequences (CTGTCTCTTATACACATCT) with command-line tool cutadapt (1.8.1), and then the trimmed files were mapped to the human genome (hg19) using Bowtie2 (2.1.0) with default parameters. Read pairs, which were aligned concordantly to the genome and had a mapping quality greater than 10, were kept for following analysis. Read pairs mapped to mitochondria DNA were discarded. Redundancy read pairs from PCR amplification were also removed afterward using Picard tools (version 1.79). The finally filtered bam files were converted into normalized TDF files using igvtools (2.26.0) for visualization of the peaks in the IGV software. Open accessible-regions for each library were defined by the peaks called by MACS2 (2.1.0) with the parameters “-g hs --nomodel --shift 0 –q 0.01”. Peaks located at blacklist genomics regions were removed using bedtools (2.25.0). These tracks shows artifact regions that tend to show artificially high signal and were identified by the ENCODE and modENCODE consortia ^87^. The filtered bam files and filtered bed files were used to generate differential peaks (FDR<0.05) using the DiffBind package in R ^88,89^. The annotations of the peaks were achieved using ChIPpeakAnno, org.Hs.eg.db, and EnsDb.Hsapiens.v75 packages in R.

### Motif analysis

For transcription factor motif analysis, the sequences of differential peaks (+/- 100 bp) were used for the motif analysis. MEME-chip^90^, as part of MEME Suite of motif-based sequence analysis tools (http://meme.nbcr.net) was used for the comprehensive motif analysis. The enriched (E value < 0.05) *de novo* DNA-binding motifs were identified by MEME-ChIP Discriminative DNA motif discovery (DREME), which uses Fisher’s Exact Test to test the significance. Transcription factors for each enriched motif were determined using Tomtom against the known motif databases (e.g. JASPAR). The set of sequences for individual matched motif were determined by Find Individual Motif Occurrences (FIMO) within the MEME suite^91^. Homer (version 4.10) was used for motif density analysis for TF of interests ^92^. Plotting of normalized tag around motif was performed using ngsplot ^93^.

### ChIP-seq and analysis

Antibody of GATA4 (Abcam, ab124265, RRID: AB_11000793) was used for ChIP-seq in this study. For each group, chromatin immunoprecipitation was performed using approximately 1×10^7^ cells. Cells were first cross-linked with 37 % formaldehyde for 10 min at room temperature, and formaldehyde was quenched for 5 min by glycine with a final concentration of 0.125 M. Chromatin was broken into small pieces with an average size of 0.5-2 kb using the Bioruptor (Diagenode). The sonicated chromatin was then incubated with 5 µg of primary antibodies overnight at 4°C. A small portion (10%) of chromatin without antibody incubation was kept as input DNA for each ChIP reaction. Subsequently, 6 g of GATA4 antibody with 75 µl of Dynabeads Protein A/G were added and incubated overnight at 4°C with overhead shaking. Magnetic beads were then washed away and chromatin was eluted. Crosslink was reversed and precipitated DNA was purified and resuspended in nuclease-free water. Sequencing libraries of immunoprecipitated DNA and input DNA were constructed according to an Illumina DNA library preparation protocol. Subsequently, ChIP-seq libraries were loaded to an Illumina HiSeq 4000 platform for deep sequencing.

The raw data were trimmed to remove the adapter sequences (including AGATCGGAAGAGCACACGTCTGAACTCCAGTCAC and AGATCGGAAGAGCG-TCGTGTAGGGAAAGAGTGTAGATCTCGGTGGTCGCCGTATCATT) with command-line tool cutadapt (1.8.1). The rest steps are the same to the ATAC-seq data analysis. Open accessible-regions for each sample were defined by subtracting input background by MACS2 (2.1.0) with the parameters “-t sample -c input”. Differential analysis was performed by DiffBind package in R ^88,89^. The annotations of the peaks were achieved using annotatePeaks.pl of the Homer software (version 4.10) ^92^. PPARGC1A-centric promoter loops were selected from promoter-capture Hi-C loops called in iPSC-derived cardiomyocytes ^45^. Processed loop calls were downloaded from ArrayExpress accession number E-MTAB-6014, and three replicate cardiomyocyte loop calls (capt-CHiCAGO_interactions-CM) were visualized using the Sushi R/Bioconductor package for genomic data visualization ^94^. Loops intersecting the GATA4 peak coordinates (chr4:23374967-23375240) are colored red.

### Network analyses

Weighted correlation network analysis (WGCNA) was used to identified gene co-expression networks ^95^. The log-transformed values of the normalized counts of all transcripts were used to perform weighted gene co-expression network analysis using WGCNA package in R. Only transcripts with a sum of values across all samples that were larger than 10 and a variance larger than 0 were used for the network analysis. For integrative network analysis, a statistical approach based on the paired expression and chromatin accessibility (PECA) was used for modeling gene regulation ^31,32^.

### Mitochondrial function and morphology analyses

Several dyes were used for investigation of mitochondrial function. For mitochondrial morphology analysis, 200 nM of MitoTracker® Green FM dye (Thermo Fisher Scientific, M7514) was used for staining live mitochondria. The images were taken using Zeiss LSM710 inverted confocal microscope, and then mitochondrial morphology was analyzed using the Fiji software ^96^. 100 nM of tetramethylrhodamine (TMRM) (Thermo Fisher Scientific, I34361) was used for measuring mitochondrial membrane potential. Fluorescence of these dyes was measured using a Tecan M1000 multimode plate reader (Tecan Systems, Inc., CA).

### Mitochondrial DNA dynamics analysis

Human mitochondrial to nuclear DNA ratio kit (Takara) was used to assess mitochondrial DNA content. Two separate primer pairs were used to generate nuclear-mitochondrial DNA content ratios. SLCO2B1 and SERPINA1 were used as nuclear genes, while ND1 and ND5 were used as mitochondrial genes. Two genes for both nuclear and mitochondrial DNA were used as an average to prevent outliers. A mathematical ratio was generated to determine the mitochondrial DNA content of each sample.

### Mitochondrial respiratory activity assay

The mitochondrial respiratory activity in cardiomyocytes was analyzed by mitochondrial stress test using a Seahorse XFp Extracellular Flux Analyzer (Agilent, CA). 45,000 of cells were plated into an XFp culture plate (Agilent) with RPMI/B27 supplemented with 10% FBS and 10 μM ROCK inhibitor. After 48h of recovering, mitochondrial stress test was performed using a Seahorse XF Cell Mito Stress Test kit (Agilent) according to the manufacture manual. Briefly, one day prior to the experiment, the XFp sensor cartridge was hydrated in XF calibrator solution and incubated overnight at 37 °C in a non-CO_2_ incubator. One hour prior to the experiment, the cells were incubated at 37 °C (non-CO_2_) in 200 μl of Seahorse assay medium, containing XF base medium supplemented 1 mM pyruvate, 2 mM glutamine, and 10 mM glucose (pH 7.4). OCR was measured with sequential injections of 2 μM oligomycin, 2 μM FCCP and each 0.5 μM of rotenone/antimycin A. Data were normalized to cell numbers for each wells, which were re-calculated using a TC20 Automated cell counter (Bio-Rad, CA) with trypan blue solution (4%)

### ATP production measurement

ATP production in differentiated cardiomyocytes were evaluated using eATP Colorimetric/Fluorometric Assay Kits (BioVision Inc, CA). Briefly, 1×10^6^ cells were homogenized in 110 μl ATP Assay buffer, and then proteins were removed using ice-cold 20 μl perchloric acid from a Deproteinizing Sample Preparation Kit (BioVision Inc, CA). After incubation on ice for 5 min, the sample were spun down at 13000 g for 2 min. Supernatant (about 120 μl) was collected and combined with 20 μl of ice-cold neutralization solution. The final solution was measured in a 96-well plate according to the protocol, and the fluorometric assay was conducted in a Tecan M1000 multimode plate reader (Excitation/Emission = 535/587 nm).

### Plasmids for of CRISPR-mediated gene activation

Two-vector system was used to establish doxycycline-inducible dCas9-VP64-mediated gene activation cells, *i.e*., lentiGuide-Puro plasmid (a gift from Feng Zhang, Addgene #52963, RRID: Addgene_137729 ^42^) expressed sgRNA and pHAGE TRE-dCas9-VP64 plasmid (a gift from Rene Maehr & Scot Wolfe, Addgene plasmid #50916, RRID: Addgene_50916 ^41^) expressed inactive version of Cas9 (dCas9) fused to a VP16 tetramer activation domain (VP64). For the sgRNA constructs, the 20-bp oligo of sgRNA was designed based on upstream of TSS region of GATA4, and then were cloned into the *BsmBI* site of the lentiGuide-Puro plasmids. The sgRNAs were designed using the on-line tool (http://crispr-era.stanford.edu) and their expression are driven by U6 promoter. The sgRNA sequences of all plasmids were confirmed by Sanger sequencing: sg_GATA4_1: GAACCCAATCGACCTCCGGC (−258bp to TSS of GATA4); sg_GATA4_2: GGTGATTCCCCGCTCCCTGG (−229bp to TSS of GATA4). Both sgRNAs were confirmed to induce GATA4 expression during early cardiomyocyte differentiation (Supplemental Figure 4A). The sg_GATA4_1 was selected in the study to investigate outcomes from overexpression of GATA4.

### Lentivirus preparation and infection in stem cells

HEK293T (ATTC, Cat# CRL-3216, RRID: CVCL_0063) cells were maintained in 6-well plates with Dulbecco’s Modified Eagle Medium (Gibco) supplemented with 10% fetal bovine serum and Penicillin Streptomycin. Packaging plasmids (pVSVg and psPAX2), lentiGuide-Puro plasmids (with sgRNAs) or TRE-dCas9-VP64 plasmids, Opti-MEM (Thermo Fisher Scientific), and X-tremeGENE 9 DNA transfection reagent (Sigma-Aldrich) were used to transfect HEK293T cells according to the manufacturer’s instructions. Media supernatant containing virus particles were filtered with 0.45μM filter and further concentrated using Lenti-X according to the manufacturers’ protocol. The stem cells were firstly infected with the pHAGE TRE-dCas9-VP64 plasmids with 300 μg/ml of G418 for selection; and then they were secondly infected with the lentiGuide-Puro plasmids with 10 μg/ml of puromycin for selection. Regarding lentiviral infection, 2 μg/ml of polybrene was used.

### Real-time QPCR

cDNA was synthesized from 1 μg of total RNA using a SuperScript VILO cDNA Synthesis Kit (Thermo Fisher Scientific) following the manufacturer’s protocol. PCR amplification was performed with a QuantStudio™ 6 Flex Real-Time PCR System (Thermo Fisher Scientific) in 20-μl reactions using 1 μl of cDNA (10 ng of total input RNA), 200 nM of each forward and reverse primer and 1X *Power* SYBR Green PCR Master Mix (Applied Biosystems). The real-time PCR program consisted of 1 cycle of 95°C for 5 min; and 40 cycles of 95°C for 15 s, 60°C 30 s and 72°C for 30 s. The β-actin was used as a normalizing gene, and relative gene expression data were calculated using the ΔΔCt method ^97^. Primers used were ordered from Qiagen, including GATA4 (QT00031997), PPARGC1A (QT00095578), and β-actin (QT00095431).

### Western blot

The cells were harvested in RIPA lysis buffer (EMD Millipore, CA) contain one tablet of Pierce™ protease and phosphatase inhibitor (Thermo Fisher Scientific), and the proteins were purified using a Branson Digital Sonifier homogenizer (Branson Ultrasonics, CT). 20 μg of protein from each sample was separated on NuPAGE 4-12% Bis-Tris protein gels (Thermo Fisher Scientific) and transferred to nitrocellulose membranes (Thermo Fisher Scientific). The protein-bound membranes were blocked with 5% of blotting-grade blocker (Bio-Rad) in PBST for one hour at room temperature and incubated with a primary antibody in 5% of blotting-grade blocker in PBST overnight at 4°C. After washing with PBST buffer, the membranes were incubated with horseradish peroxidase (HRP)-conjugated-secondary antibody for 1h at room temperature. The membranes were developed with SuperSignal West Femto Maximum Sensitivity Substrate (Thermo Fisher Scientific) and exposed on a ChemiDoc Touch imaging system (Bio-Rad) for imaging. The primary antibody used in this study included mouse anti-GATA4 (R&D systems, MAB2606, RRID: AB_2108599), rabbit anti-PGC1 alpha antibody (Abcam, ab54481, RRID: AB_881987), rabbit anti-HA-tag (Cell Signaling Technology, #5017, RRID: AB_10693385), and mouse anti-beta-actin (Thermo Fisher Scientific, MA5-15739, RRID: AB_10979409). The secondary antibodies include HRP-conjugated-horse anti-mouse IgG (Cell Signaling Technology, 7076S, RRID: AB_330924) and HRP-conjugated-goat anti-rabbit IgG (Thermo Fisher Scientific, 32460, RRID: AB_1185567).

### Maternal TKI exposure in mice model

The Stanford Institute of Medicine Animal Care and Use Committee approved all protocols. 6-week-old female C57BL/6 mice purchased from Jackson Laboratory (Bar Harbor, ME) were used. Female and male were housed as a 1:1 ratio in a single cage in the afternoon (∼4 pm). In the following morning (designated as E0.5), females with vaginal plugs were injected with sunitinib and saline as control. TKIs or saline were injected intraperitoneally every other day towards E18.5. The doses for each drug per female mouse (∼20 g) per injection (in 200 µL saline) was as follows: sunitinib and vandetanib, 100 µg; Imatinib, 1.5 mg. In the morning of E19.5, the plasma from newborns (P0) and their mom were collected by the following protocol. Blood samples were collected directly into EDTA-treated tubes. The tubes were shaken gently, but thoroughly afterward. Samples were then centrifuged at 20-24 °C at 4500 g for 10 min. Then the plasma was transferred and aliquoted into pre-cooled Eppendorf tubes without aspirating blood cells. Plasma samples were frozen immediately on dry ice and stored at -80 °C.

### Detection of Drug Exposure by LC-MS

The detailed methods used for metabolomic sample preparation, data acquisition, and analysis have been described previously ^98^. Briefly, plasma metabolites were extracted with organic solvent for LC-MS analysis. Samples were separated using RPLC (Zorbax-SB-aq column 2.1 x 50mm, 1.7mm, 100Å; Agilent) and collected in positive ion mode on a Thermo Q Exactive HF mass spectrometer (Thermo Fischer Scientific). Raw data was processed with Progenesis QI 2.3 software (Water, Milford, MA, USA) to align and quantify chromatographic peaks. Imatinib (m/z: 494.2663) and sunitinib (m/z: 399.2191) were identified by accurate mass, retention time, MS/MS fragmentation, and quantified relative to standard curves.

### Statistical analysis

Statistical analysis was performed using GraphPad Prism 8.4 (GraphPad Software, Inc., San Diego, CA, RRID: SCR_002798). Nonparametric T-test was used to compare data between two groups. Data are reported as means ± standard error of the mean (SEM).

### Data Availability

The RNA-seq, ATAC-seq, and ChIP-seq data generated for this work have been deposited in NCBI Gene Expression Omnibus, and they are accessible numbers are GSE149586 for RNA-seq, GSE149589 for ATAC-seq, and GSE149591 for ChIP-seq.

## SUPPLEMENTAL FIGURES & TABLES

**Supplemental Figure 1. No significant differences in efficiency of CM differentiation were observed upon exposure to NOAELs of drugs. (A)** Determination of the no-observed-adverse-effect-levels (NOAELs) of drugs used in this study. **(B)** Flow cytometry analysis differentiated cardiomyocytes of each group. The differentiated cardiomyocytes (day20) were labeled with TNNT2 antibodies and used for the flow cytometry analysis. The data were collected from Exposure II, *i.e*., the cells were exposure to TKI drugs till the differentiated CMs were collected for analyses.

**Supplemental Figure 2. Evaluation of Ca**^**2+**^**-handling of differentiated cardiomyocytes and expression of genes related to contractile. (A)** Bar charts represent developmental exposure to NOAELs of immunosuppressant (tacrolimus and mycophenolate) and thalidomide caused alternations of Ca^2+^- handling properties, including decreases in beating rate and amplitude, and increased in time to peak, TD90/50, and decay tau. The data were collected from Exposure II. The *p* values were calculated by nonparametric T-test between control and TKI-treated groups. NA, no significance was observed (*p*>0.05). **(B)** Gene expression of genes related to contraction of cardiomyocyte. The color of the heatmap represents log2-transformed fold changes, which was calculated by treatment vs. DMSO using the normalized counts by trimmed mean of M-values (TMM) normalization method.

**Supplemental Figure 3. Motif analysis in TKI-treated cells and control cells from ATAC-seq data. (A)** Enriched de novo motifs for MEF2s family were discovered using MEME-chip based on down-regulated chromatin accessibility in TKI-treated cells. **(B)** Peak signals (+/- 1000 bps) arounds submits of the motifs for GATA4, TEAD1, and TEAD3 in TKI-treated cells compared with DMSO-treated cells. Binding intensities are shown as sequencing depth normalized tag count.

**Supplemental Figure 4. Gene expression of GATA4 during cardiomyocyte differentiation. (A)** Data represent normalized counts collected from three time points of the RNA-seq data: day 0, day 6, and day 20. **(B)** Protein expression analysis of GATA4 in two sgRNA designs. Two sgRNA were designed for targeting upstream of TSS of GATA4. Cells from intermediate differentiation stages were collected for protein analysis using Western blot

**Supplemental Figure 5. Changes of mitochondrial DNA copies during cardiomyocyte differentiation upon TKI exposure. (A)** Dynamic mitochondrial DNA (mtDNA) copies during cardiomyocyte differentiation in control (0.1% DMSO). Data were collected from four time points: day 6, day 9, day 23, day 26. (**B-C)** TKI-exposure decreased mtDNA copies compared to control. However, no significant differences were observed. NA, *p*>0.05. The *p* values were calculated by nonparametric T-test between control and TKI-treated groups. **(D)** Overexpression of GATA4 by adding doxycycline (dox) increased mtDNA copies in TKI-treated cells. Data were collected from cardiomyocytes of day26. The *p* values were calculated between TKI-treated cells with and without overexpressing GATA4. **(E)** Protein expression analysis of PGC-1α in cardiomyocytes.

**Supplemental Figure 6. Maternal exposure to sunitinib induced moderate pathology in offspring. (A)** Experimental design of maternal exposure using mouse model. **(B)** Example of death of fetus due to maternal exposure to sunitinib. **(C)** Moderate CHD-like histopathologic outcomes were observed in fetal mice (E14.5). TM, thinner myocardium; VSD, ventricular septal defect. Scale bars: 200 µm. **(D)** Bioaccumulation of sunitinib was found in the blood of both female mouse and newborns (P0). Only one adult female mouse was used.

**Supplemental Table 1. Gene lists of each module from WGCNA.**

**Supplemental Table 2. Summary of each module from WGCNA.**

**Supplemental Table 3. ChIP-seq analysis of GATA4 occupancy in cardiomyocytes from both control and sunitinib-treated groups.**

## ACKNOWLEDGEMENTS

We thank both the Stanford Cardiovascular Institute (SCVI) Biobank and Stem Cell Core Facility of Genetics, Stanford University, which provided the human pluripotent cells. We thank Ningyi Shao, Joe Zhang and Ning Ma of SCVI for their advice in stem cell experiments. We thank Prof. Ronald Davis of Stanford University for supporting XFp Seahorse analyzer. This work used the Genome Sequencing Service Center by Stanford Center for Genomics and Personalized Medicine Sequencing Center, supported by the grant award NIH S10OD020141. We thank Stanford Neuroscience Microscopy Service (NMS, supported by NIH NS069375) for providing the confocal microscope. This work was support by California Institute for Regenerative Medicine, CIRM GC1R-06673-A (M.P.S.), American Heart Association Career Development Award (18CDA34110128 to Q.L. and 18CDA34110293 to M-T.Z.), Stanford Child Health Research Institute and the Stanford NIH-NCATS-CTSA, UL1 TR001085 (Q.L. and M-T.Z.), NIH Pathway to Independence Award (HL133473-01A1 to H.W. and K99HG010362 to K.V.B.), NIH R01 HL141851, NIH R01 HL113006, and NIH R01 HL141371 (J.C.W.), NIH and a European Research Council Advanced Investigator Grant and Steinmetz Cardiomyopathy Fund (L.M.S.). The contents of this publication are solely the responsibility of the authors and do not necessarily represent the official views of CIRM or any other agency of the State of California.

## AUTHOR CONTRIBUTIONS

Q.L. conceived and designed the experiments under the supervision of M.P.S.. Q.L. performed most of experiments and data analysis. H.W performed Ca^2+^ handling analysis and microscopy. Q.J.L. and J.L. performed the mice studies. C.J. and Z.D. performed network analysis. M.T.Z. carried out the flow cytometry and analysis. D.P.M. and B.L-M. carried mass spectrometry. K.V.B, C.Z., N.K.W., and L.T. provided assistance for bioinformatics. B.Z., A.M.N., J.J.G., A.M.L., H.G., M.S.T, T.F., E.W., and Y.L. provided helpful assistance. L.M.S., W.H.W., M.A.K., and J.C.W. provided valuable insights and helpful assistance. Q.L. and M.P.S. wrote the manuscript with input from all authors.

