## Supplementary material for "Tyrosine kinase inhibitors induce mitochondrial dysfunction during cardiomyocyte differentiation through alteration of GATA4-mediated networks": Supple.Figures_and_tables: Supplemental Figures1-6.pdf

### Supplemental Figure 1

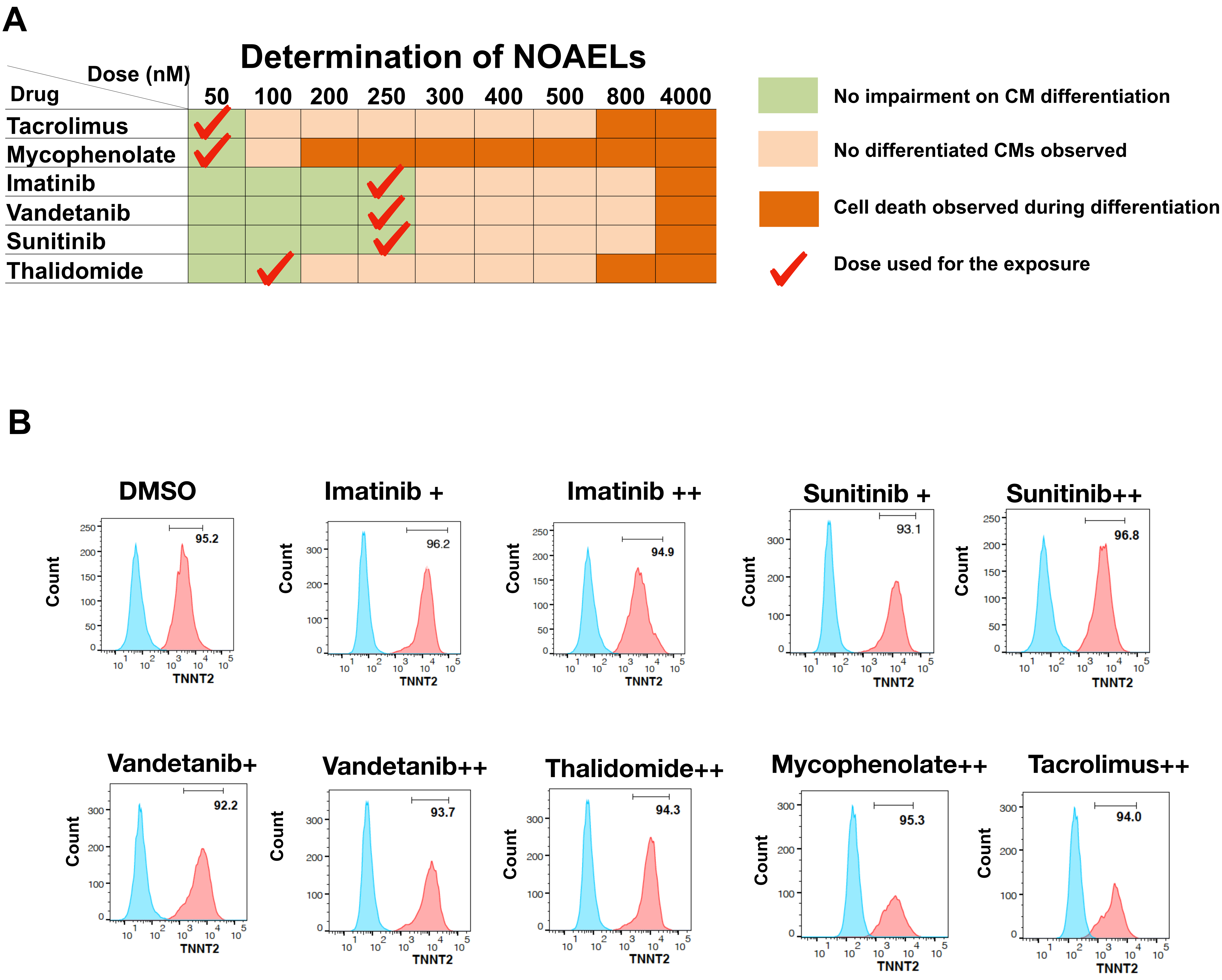

### Supplemental Figure 2

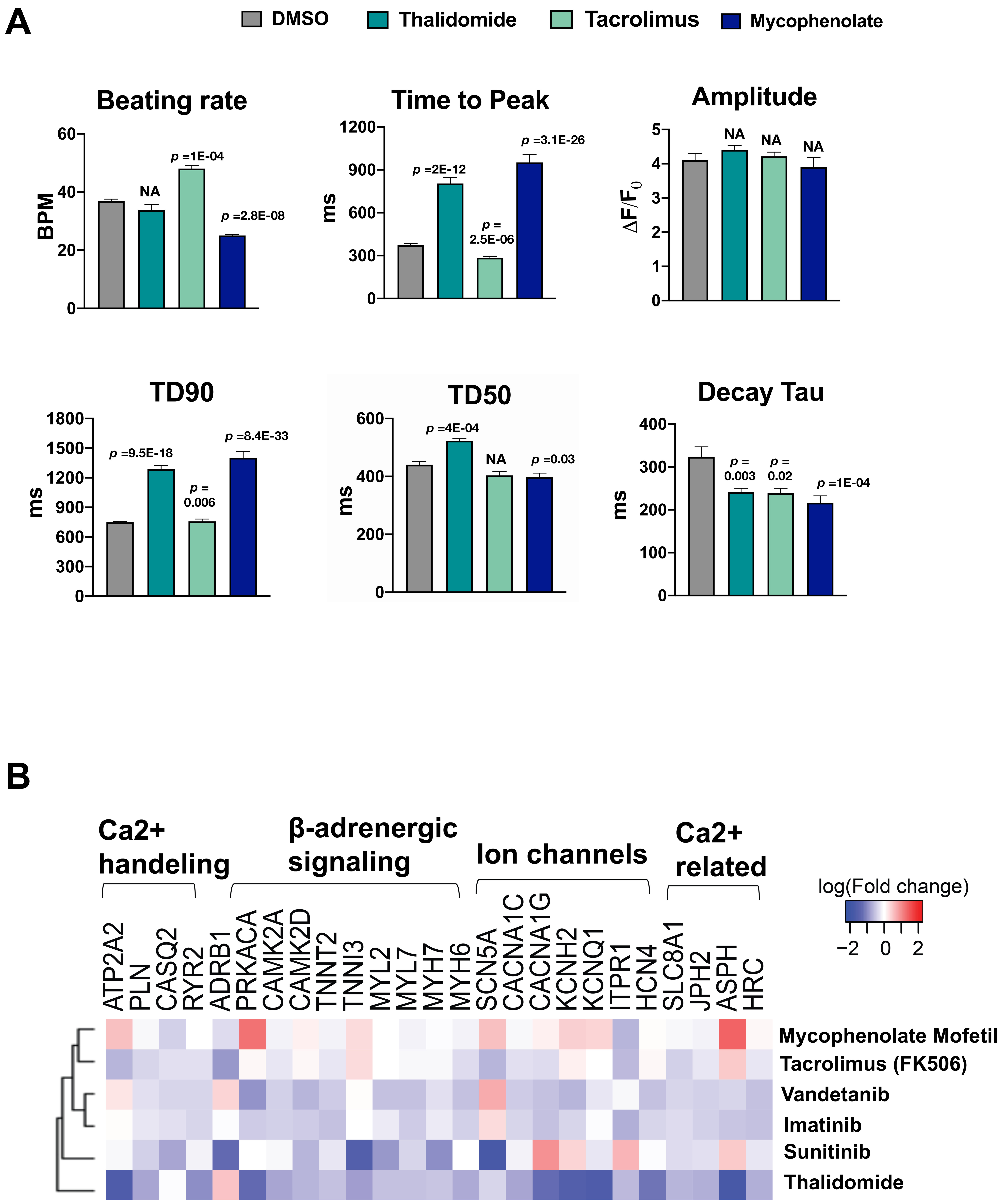

### Supplemental Figure 3

**A**

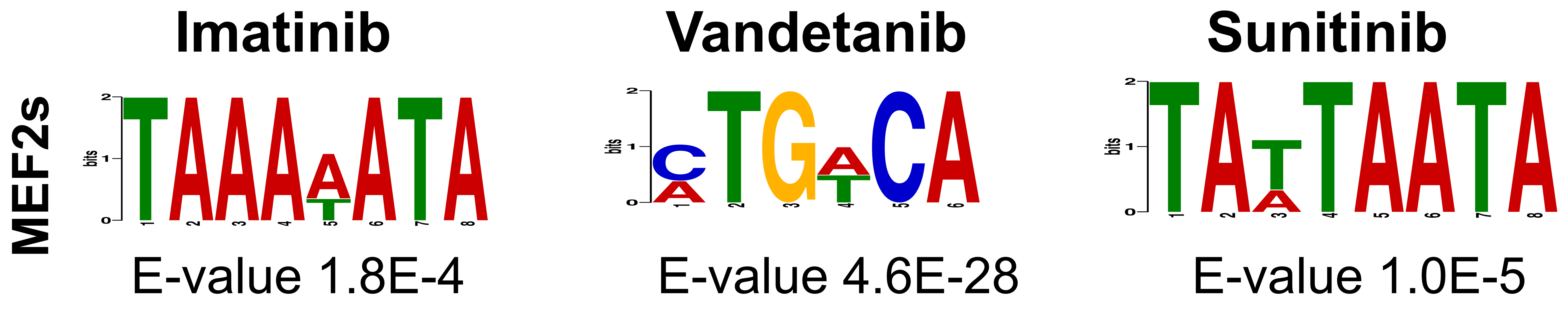


**B**

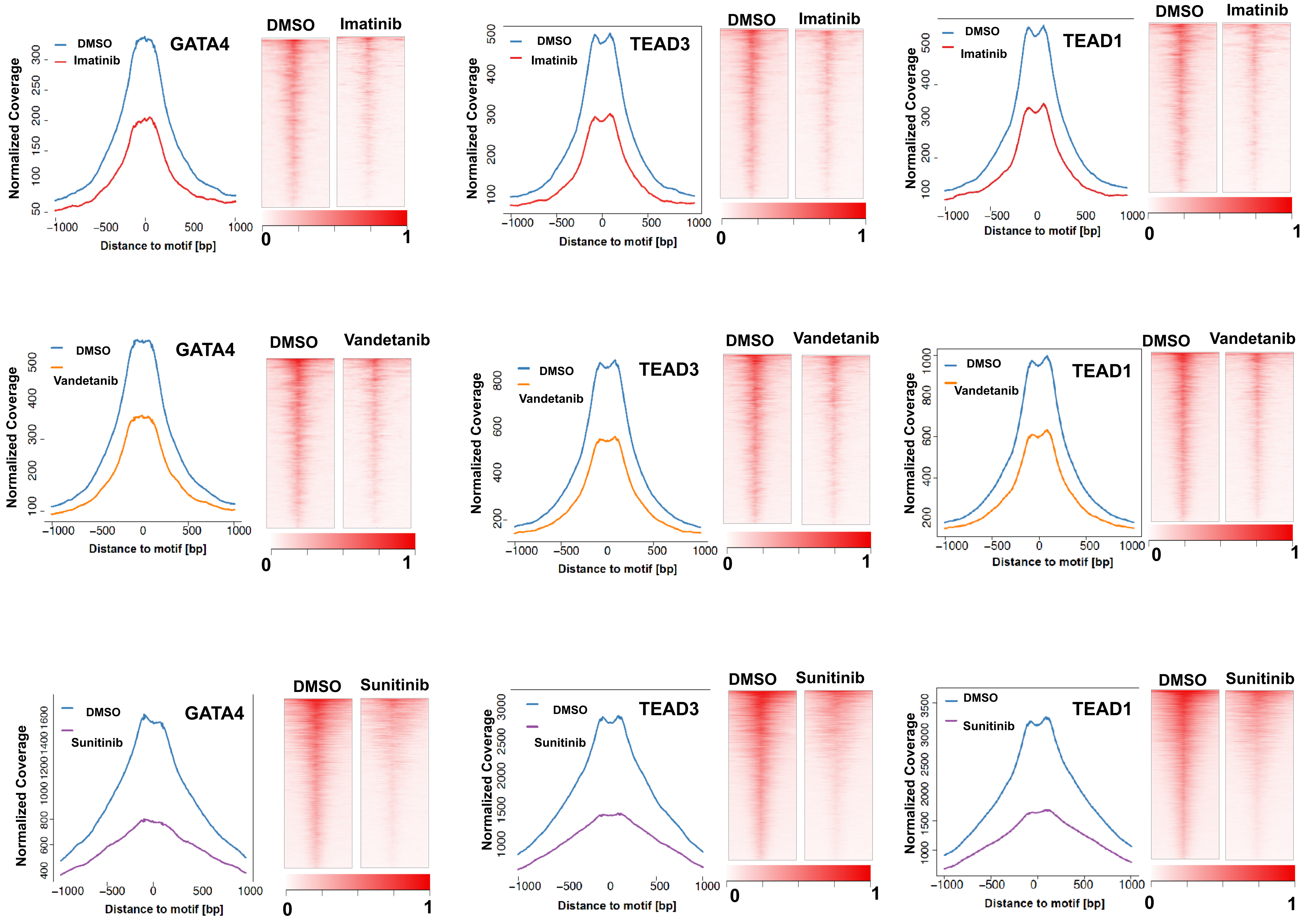

### Supplemental Figure 4

A

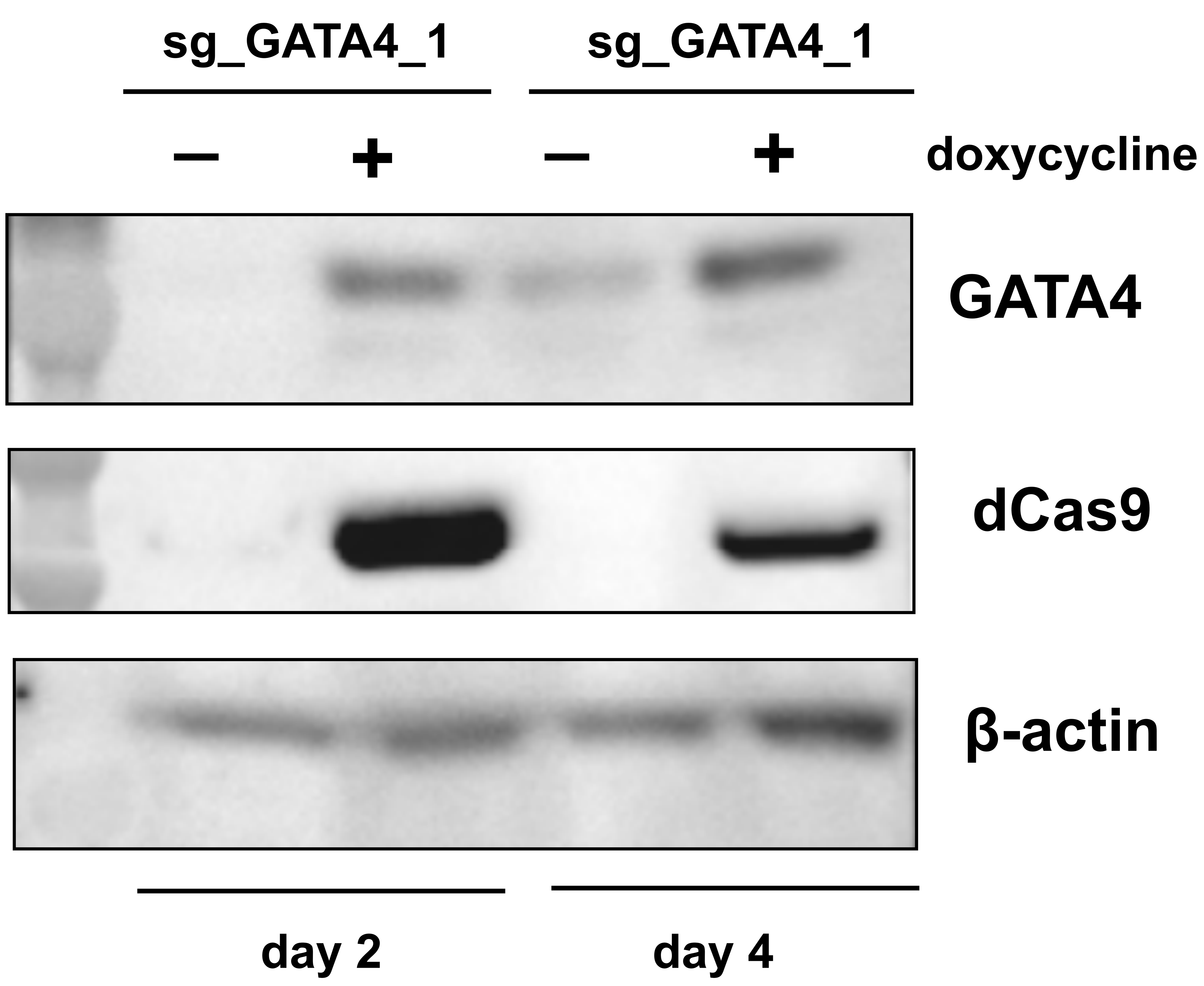


B

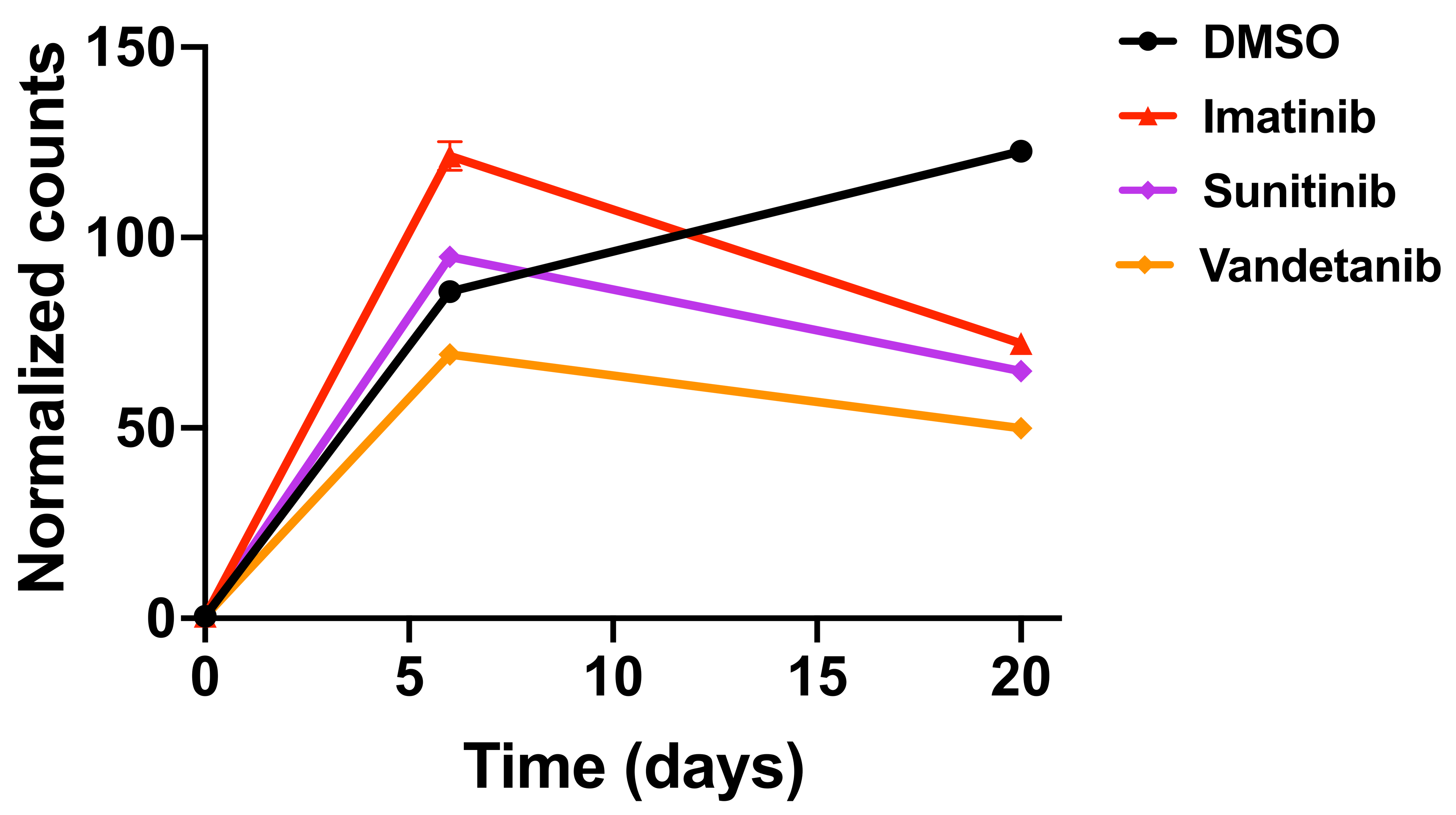

### Supplemental Figure 5

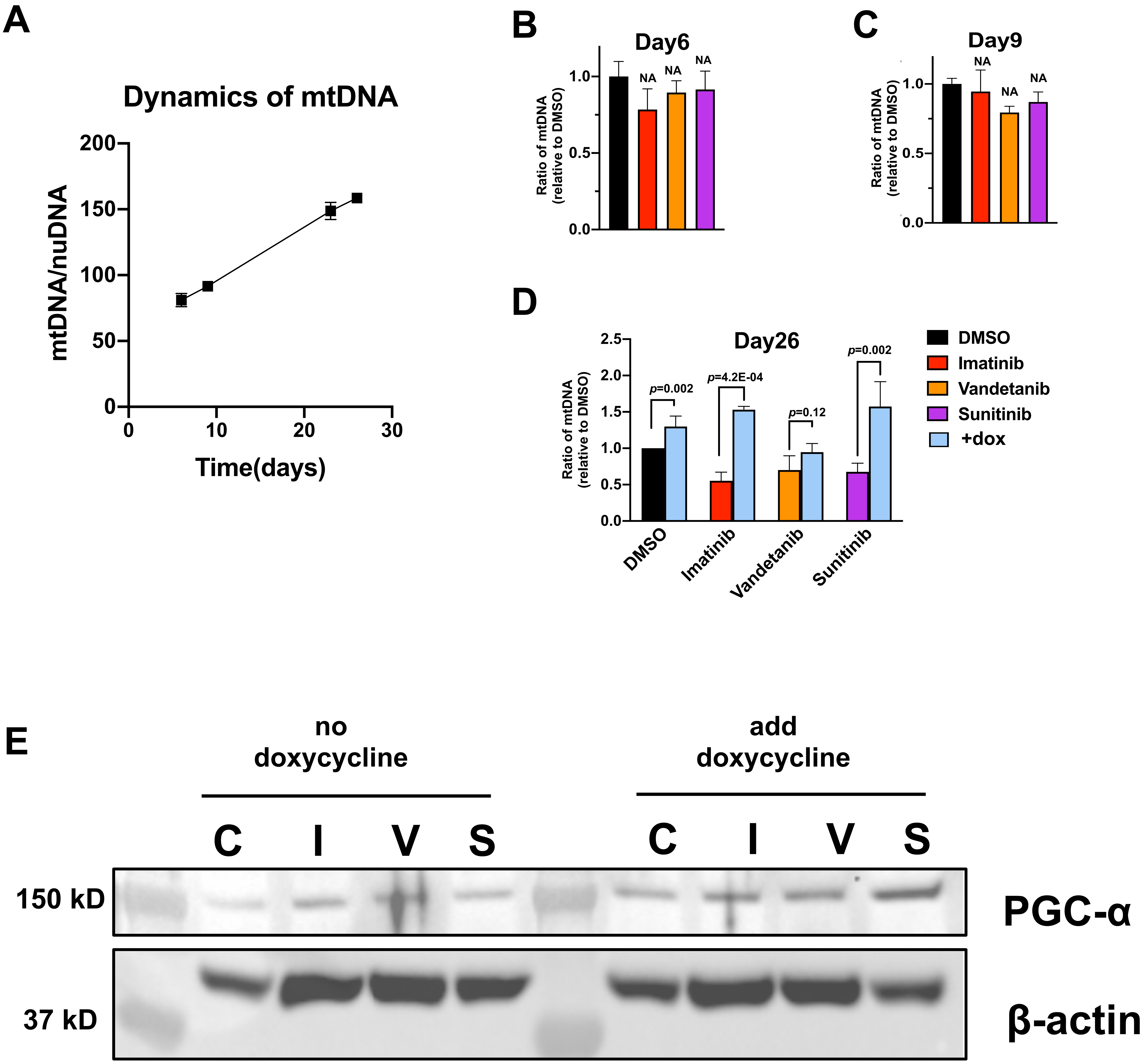

### Supplemental Figure 6

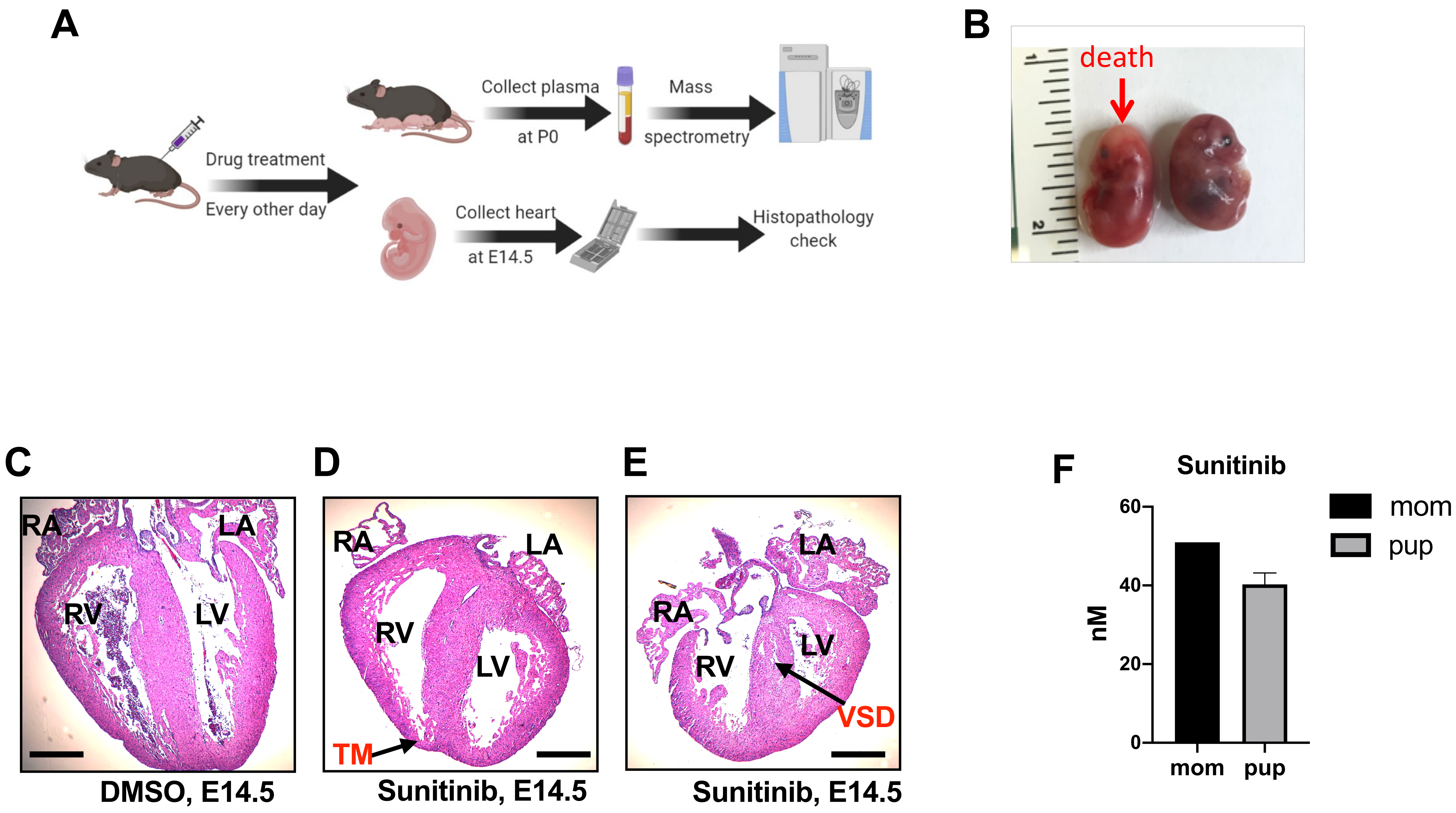
